# Syndrome-informed phenotyping identifies a polygenic background for achondroplasia-like facial variation in the general population

**DOI:** 10.1101/2023.12.07.570544

**Authors:** Michiel Vanneste, Hanne Hoskens, Seppe Goovaerts, Harold Matthews, Jose D Aponte, Joanne Cole, Mark Shriver, Mary L. Marazita, Seth M. Weinberg, Susan Walsh, Stephen Richmond, Ophir D Klein, Richard A Spritz, Hilde Peeters, Benedikt Hallgrímsson, Peter Claes

**Affiliations:** Department of Human Genetics, KU Leuven, Leuven, Belgium; Department of Cell Biology & Anatomy, Cumming School of Medicine, University of Calgary, Calgary, AB, Canada; Alberta Children’s Hospital Research Institute, Cumming School of Medicine, University of Calgary, Calgary, AB, Canada; McCaig Bone and Joint Institute, Cumming School of Medicine, University of Calgary, Calgary, AB, Canada; Medical Imaging Research Center, University Hospitals Leuven, Leuven, Belgium; Department of Biomedical Informatics, University of Colorado School of Medicine, Aurora, CO, USA; Department of Anthropology, Pennsylvania State University, State College, PA, USA; Center for Craniofacial and Dental Genetics, Department of Oral and Craniofacial Sciences, School of Dental Medicine, University of Pittsburgh, Pittsburgh, PA, USA; Department of Human Genetics, School of Public Health, University of Pittsburgh, Pittsburgh, PA, USA; Department of Biology, Indiana University Purdue University Indianapolis, Indianapolis, IN, USA; Applied Clinical Research and Public Health, School of Dentistry, Cardiff University, Cardiff, UK; Department of Orofacial Sciences and Program in Craniofacial Biology, University of California, San Francisco, CA, 94143, USA; Department of Pediatrics, Cedars-Sinai Medical Center, Los Angeles, CA 90048, USA; Department of Pediatrics, University of Colorado School of Medicine, Aurora, CO, USA; Department of Electrical Engineering, ESAT/PSI, KU Leuven, Leuven, Belgium

**Keywords:** Craniofacial Variation, Developmental Constraints, Complex Traits, Mendelian Disease, Achondroplasia, Genome-Wide Association Study

## Abstract

Human craniofacial shape is highly variable yet highly heritable with genetic variants interacting through multiple layers of development. Here, we hypothesize that Mendelian phenotypes represent the extremes of a phenotypic spectrum and, using achondroplasia as an example, we introduce a syndrome-informed phenotyping approach to identify genomic loci associated with achondroplasia-like facial variation in the normal population. We compared three-dimensional facial scans from 43 individuals with achondroplasia and 8246 controls to calculate achondroplasia-like facial scores. Multivariate GWAS of the control scores revealed a polygenic basis for normal facial variation along an achondroplasia-specific shape axis, identifying genes primarily involved in skeletal development. Jointly modeling these genes in two independent control samples showed craniofacial effects approximating the characteristic achondroplasia phenotype. These findings suggest that both complex and Mendelian genetic variation act on the same developmentally determined axes of facial variation, providing new insights into the genetic intersection of complex traits and Mendelian disorders.

## Introduction

Genetic variation in conjunction with environmental factors influences developmental processes that drive phenotypic variation^1,2^. Rare major-effect variants and common variants have been identified through largely separate studies of monogenic and complex phenotypes, respectively^3^. However, recent advances have led to a far deeper understanding of the relationship between normal and syndromic development. One key conceptual hypothesis is that both normal and syndromic phenotypic variation occur predominantly along developmentally delimited directions of phenotypic change, or ‘axes of variation’. Both common and rare variants can act upon these axes, causing syndromic phenotypic variation to occur along the extremes of normal phenotypic axes of variation. Preliminary findings supporting this^3–8^ highlight the potential importance of an integrated approach that incorporates both complex and Mendelian traits into the study of phenotypic variation.

Human facial shape shows great potential for such an integrated approach. Facial shape is an assemblage of highly variable, developmentally complex phenotypes that are largely genetically determined, involving both common and rare variants with a range of effect sizes^9^. Recent advances in three-dimensional (3D) image processing technology and genome-wide association studies (GWAS) have enabled the identification of hundreds of genetic loci associated with normal-range facial variation, yet collectively these only account for about 10% of facial phenotypic variance^9^. In addition, many rare large-effect variants have been discovered through the study of Mendelian disorders with craniofacial dysmorphism^10^. A well-known example is the recurrent pathogenic gain-of-function variant G380R in *FGFR3* that causes achondroplasia (ACH), the most common form of skeletal dysplasia^11^. FGFR3 is a regulator of bone growth that is expressed in chondrocytes and mature osteoblasts, and increased FGFR3 signaling suppresses proliferation and maturation of growth plate chondrocytes. This, in turn, impairs endochondral bone growth, resulting in rhizomelic limb shortening and short stature in ACH^12,13^. In the skull, premature fusion of skull base synchondroses leads to a shortened basicranium and a recognizable pattern of frontal bossing and midface hypoplasia in affected individuals^14^. While all ACH patients share the same pathogenic G380R *FGFR3* variant, a modest range of variability exists within the characteristic ACH phenotype^15^. As for most monogenic disorders, the factors underlying this variable phenotypic expressivity remain largely unknown.

If both normal-range and syndromic phenotypic effects converge on developmentally delimited axes, and phenotypic variation occurs principally along those axes, we would expect that facial variation along a syndromic phenotypic axis would also be present to some degree in the general population. In this work, we tested this hypothesis by projecting the ACH facial phenotype onto an unselected control population to model ACH-derived facial variation as a quantitative trait, rather than a binary categorical (or monogenic) trait. Genetically mapping these traits in the control population revealed strong enrichment for genes involved in developmental processes that are key in the pathophysiology of ACH. Elements of the ACH phenotype could also be replicated in silico in two independent control samples, relying solely on the uncovered polygenic background. We discuss the implications of our findings to the broader field of Mendelian and complex trait genetics.

## Results

### ACH phenotype can be constructed from axes of normal-range facial variation

3D facial photographs were available from 43 subjects with ACH and 8246 unselected controls, all with European ancestry (Fig. 1a). We obtained homologous facial configurations by non-rigidly mapping an atlas (composed of 7160 points) to each individual image^16^. Controls were Procrustes aligned to a common coordinate system and principal component analysis (PCA) was applied to capture the major axes of normal-range facial variation. Projecting ACH subjects into the same coordinate space accurately described syndromic facial shape variation, with 0.49 mm average error between the original ACH shapes and corresponding projections (Supplementary Fig. 1a). Regions of higher reconstruction error coincided with the clinically most distinct regions (e.g., nasion and forehead). As a comparison, the average PCA reconstruction error of the controls was 0.35 mm (Supplementary Fig. 1b). While ACH syndromic samples could be coded well as linear combinations of these axes (principal components) of normal variation, they showed greater variation overall and were generally found towards the tail-end of the distribution (Supplementary Data 1).

**Figure 1.**
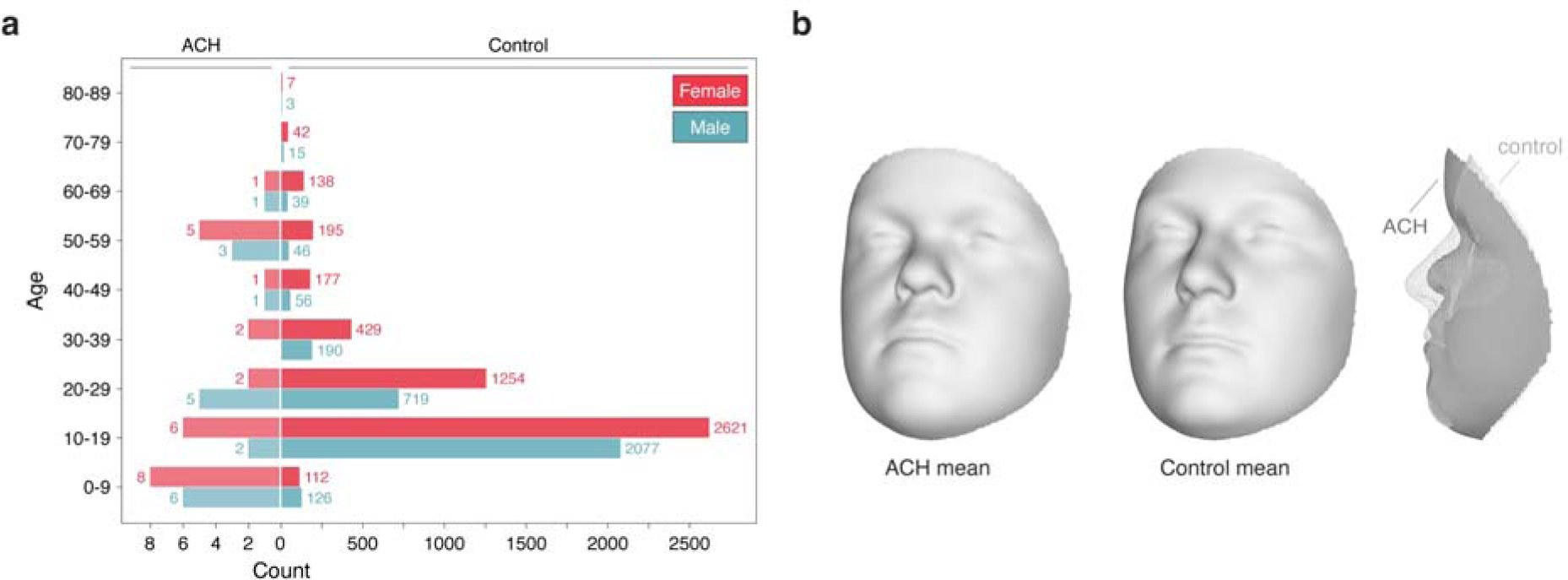
Sample characteristics of the achondroplasia and control dataset. (a) Age and sex distribution, (b) Average facial shapes.

### Definition of ACH-informed phenotype as a quantitative trait

We compared 3D facial images of the ACH samples to the unselected controls (Fig. 1b) at multiple scales, starting from a global description of facial shape and gradually focusing on more local segments of shape variation (Supplementary Fig. 2a) determined by hierarchical spectral clustering^17^. By regressing facial shape onto syndrome status (ACH or control), we found that facial shape was significantly different between ACH and control samples in 58 out of the 63 facial segments (p < 0.05) (Supplementary Fig. 2b). For each of these significant segments, we established an ACH trait axis as the vector spanning the ACH and control shape means. These axes describe the facial shape effects associated with ACH (“ACH-derived facial trait”), such as frontal bossing and midfacial hypoplasia. Moving along the axes is equivalent to changing phenotypic severity (Fig. 2a).

**Figure 2.**
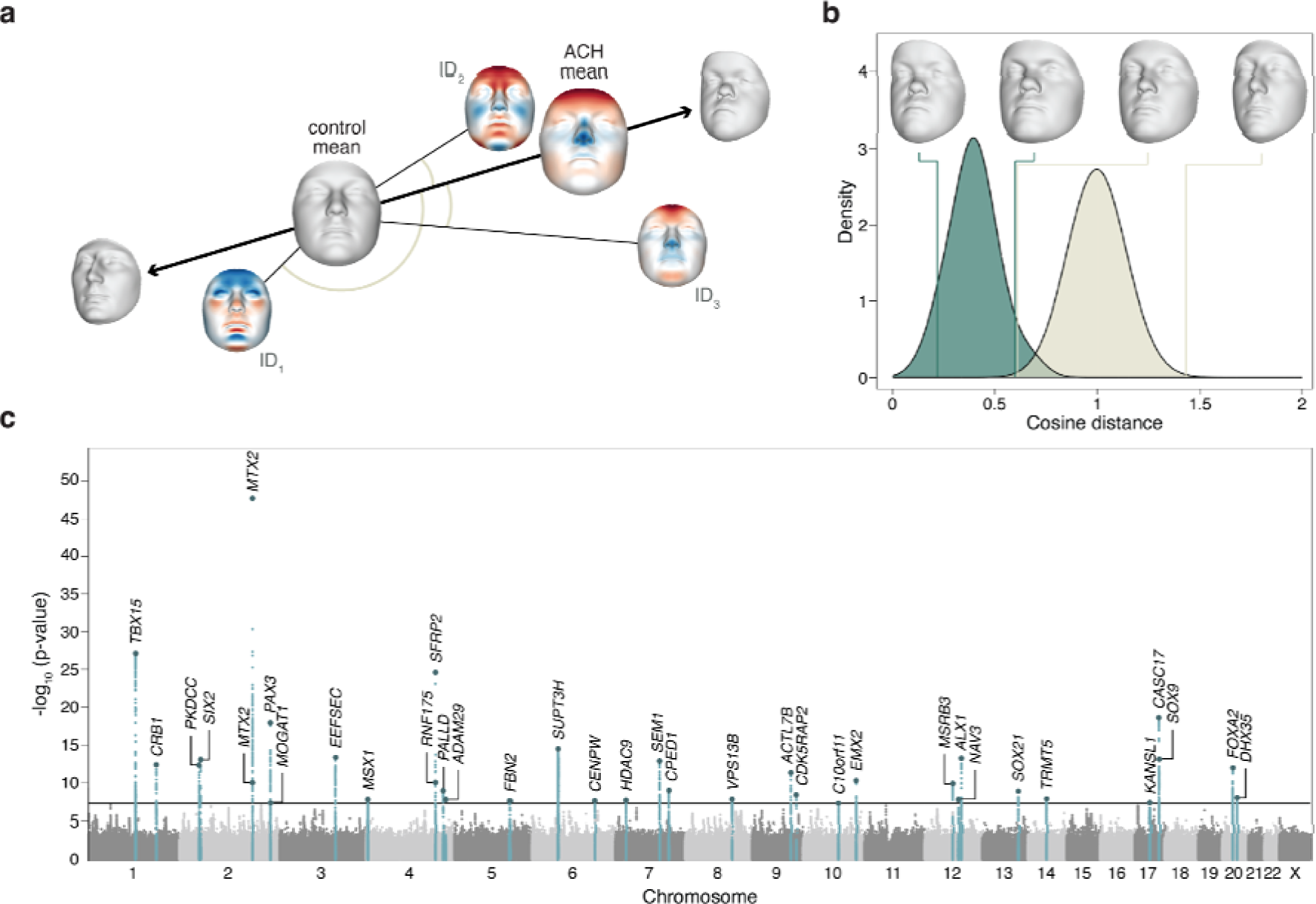
Achondroplasia-informed phenotyping. **(a)** ACH trait axis spanning the ACH and control mean shapes. Morphs on the left and right side of the axis represent the extremes of the phenotypic spectrum. Controls (ID_1-3_) can be scored along the axis by measuring the angle between their individual vectors and the ACH trait vector. Facial variation of the three control individuals is visualized as a heatmap. Red areas on the facial shape correspond to a local outward deviation from the control mean shape, blue indicates inward deviation. **(b)** Distribution of the facial trait scores for the full face (segment 1) for both the ACH (in green) and control (in beige) datasets. Values smaller than 1 indicate more ACH-like; values greater than 1 indicate less ACH-like. The mean facial shape of the 5 lowest and highest scoring individuals is shown for both ACH and control samples. **(c)** Manhattan plot of genomewide associations. For each SNP, the lowest p-value (CCA, right-tailed chi square) across all 58 significant facial segments is plotted. The full horizontal line represents the genome-wide significance threshold (p = 5e-8). Candidate genes are annotated to each genome-wide significant locus (n = 35).

We quantified the extent to which the ACH-derived traits or shape effects exist in the control population by measuring facial similarity of the unselected controls to the ACH trait axes using the cosine distance, hereafter referred to as “syndrome-informed phenotyping” (Fig. 2a). This approach generated univariate scores, with controls that display ACH-like facial features having a low score (values smaller than 1), while individuals with an inverse phenotype (e.g., protrusion of the midface) have higher scores (values greater than 1) (Fig. 2b). The control sample showed great variation in the ACH trait scores, yet a clear overlap was observed with the ACH cohort scored along the same axes (Supplementary Data 2). Furthermore, the ACH trait scores explained 2.6% of full facial shape variation in the control population, which is substantial, considering that sex and age explained 11.3% and 4.9% of variance in the same cohort, respectively.

### Multivariate GWAS reveals polygenic background of ACH-derived facial traits

We sought to identify SNPs associated with facial variation in the general population along the ACH trait axes. We combined the trait scores for all 58 significant facial segments into a matrix, for the US and UK subsamples separately, and performed a multivariate GWAS meta-analysis using canonical correlation analysis. In total, we identified 1925 SNPs that reached genome-wide significance (p <15e−8). Significant SNPs were merged into 35 genomic loci, revealing a polygenic basis for normal facial variation along a characteristic shape axis derived from ACH, a monogenic disorder (Fig. 2c; Supplementary Table 1). The 35 lead SNPs combined explained 1.77% and 2.00% of total facial shape variation in the US and UK subsamples, respectively. While some lead SNPs correlated most with ACH-derived shape changes of the full face, others showed more localized effects, affecting only a specific aspect of the ACH facial phenotype (Supplementary Data 3).

For many of these significant associated loci, candidate genes in the immediate vicinity have well-established roles in craniofacial development and/or have previously been identified in GWAS of facial shape variation (Supplementary Table 1). To our knowledge, 4 of the 35 loci have not previously been linked with craniofacial morphology. No significant associations were found for SNPs near *FGFR3*; neither did we find significant enrichment for associations with genes that interact with *FGFR3* directly (p = 0.48) (Supplementary Table 2). However, STRING analysis of the GWAS-associated candidate genes showed plausible interactions with the FGFR3 network at higher levels (Supplementary Fig. 3).

### Genetic loci associated with ACH-derived facial variation are enriched for processes related to skeletal development

Gene-set enrichment analysis^18^ of the 35 associated loci showed significant enrichment for biological processes related to cartilage growth and development, and skeletal development overall (Supplementary Table 3). To evaluate targeted enrichment of certain biological processes, we compared our findings to those of a GWAS of normal facial variation performed in the same unselected control group by White et al.^19^. While the normal facial GWAS showed enrichment for a broad spectrum of processes related to embryonic development, the current ACH-informed GWAS was enriched for a specific subset of these biological processes, with all but one term (84/85, 99%) also significantly enriched in the previous study^19^. Processes related to cartilage development such as chondrocyte differentiation and development, chondrocyte hypertrophy, and cartilage condensation were consistently more enriched in the ACH-informed GWAS than in the previous uninformed facial shape GWAS by White et al.^19^ (Fig. 3a). Interestingly, these same biological processes are at the core of ACH pathophysiology^20^. For other branches of system development (e.g., nervous system development, circulatory system development), we observed no significant difference between the ACH-informed and uninformed facial shape GWAS^19^ (Fig. 3a). Similarly, the ACH-informed GWAS genes were specifically enriched for skeletal developmental genes when compared against all genes previously identified through GWAS of facial shape (Fig. 3b), as well as against known craniofacial genes implicated in Mendelian syndromes^9^ (Supplementary. Fig 4a-b; Supplementary Table 4). A similar targeted enrichment was not observed in comparison to a “negative control” GWAS of inflammatory bowel disease^21^ (Supplementary Fig. 4c).

**Figure 3.**
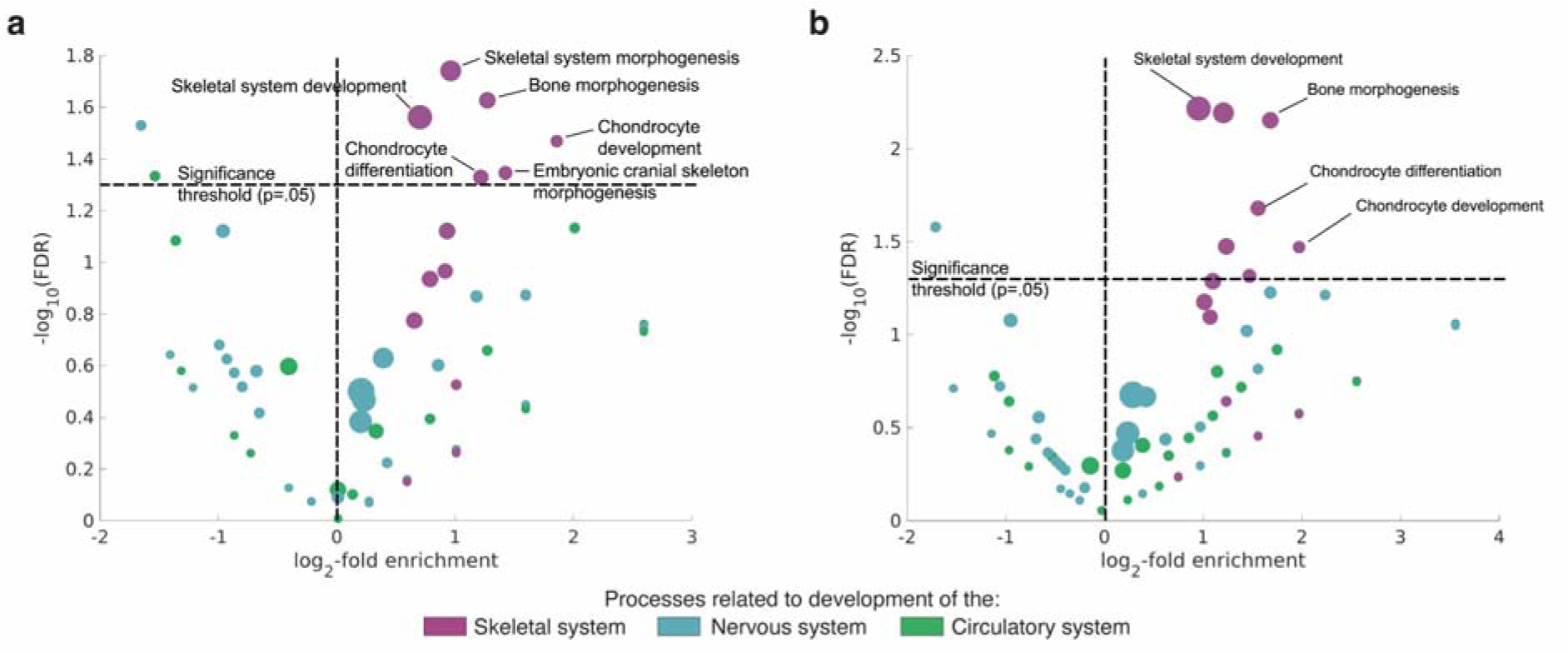
GO enrichment analysis. Fold enrichment of Gene Ontology (GO) biological processes enriched in the ACH GWAS compared to different background sets. **(a)** ACH-informed GWAS versus uninformed GWAS of normal facial variation by White et al. **(b)** ACH-informed GWAS versus all genes previously identified through GWAS of facial shape. Only processes enriched in both studies are displayed. Node size corresponds to the number of genes mapped to each process.

**Figure 4.**
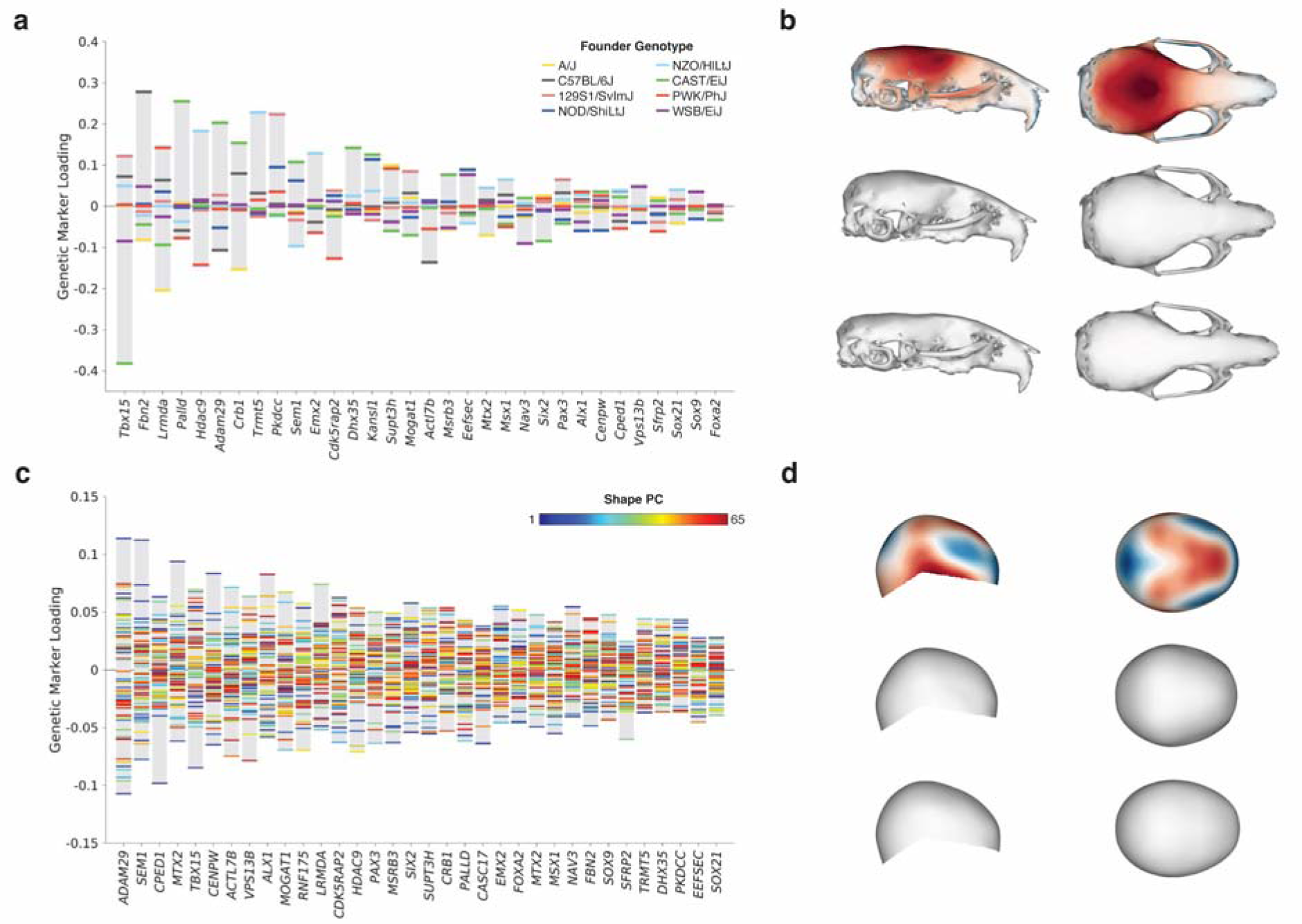
Multivariate genotype-phenotype mapping of mouse and human craniofacial shape. Genetic marker loadings for the multivariate genotype-phenotype mapping of the GWAS candidate genes onto **(a)** mouse craniofacial shape and **(c)** human cranial vault shape. Genes are ordered by their relative contribution to the associated shape effects shown in **(b)** and **(d)**, respectively. The top row shows the mean craniofacial shape colored according to the difference between the upper and lower extremes of the MGP shape axis. Red areas indicate a local inward deviation, blue indicates an outward deviation. The middle row shows the upper extreme of the MGP shape axis. The bottom row shows the lower extreme of the same shape axis.

### Genetic loci associated with ACH-derived facial trait are not enriched for layer-specific murine growth plate gene expression

Following the observed enrichment for cartilage-related processes, we further tested whether the ACH-informed GWAS genes were preferentially expressed in certain epiphyseal layers or at specific chondrocyte maturation stages. We analyzed public gene expression data from murine growth plates and chondrocytes, but found no correlation between MAGMA gene-level p-values and gene expression specificity per epiphyseal layer or chondrocyte maturation stage. Differential expression analysis showed no significant associations between gene-level p-values and changes in gene expression between early and late chondrocyte maturation stage.

### ACH-derived facial trait shows significant genetic correlations with other ACH-linked traits

We calculated the Spearman genetic correlation^22^ between the ACH-derived facial trait and five ACH-associated traits, including body height, infant head circumference, obstructive sleep apnea, lung volume, and sitting height ratio (Supplementary Table 5). Although these five traits are all associated with the pathogenic G380R *FGFR3* variant in individuals with ACH, it is unknown if they also correlate with ACH-like facial features in the healthy population. We found significant genetic correlations (FDR-corrected p < 0.05) between the ACH-derived facial trait and body height, obstructive sleep apnea, and sitting height ratio (Table 1), likely pointing to the same skeletal system pathways that showed enrichment in the previous analysis. In individuals with ACH, reduced endochondral bone growth causes disproportionate short stature with increased sitting height ratio, and can also lead to narrowing of the upper airway, which in turn may increase the risk for obstructive sleep apnea^23,24^. The ACH-derived facial trait also showed a stronger genetic correlation with sitting height ratio than with height, while uninformed facial shape by White et al.^19^ showed no differential correlation with these two traits. In line with the current findings, previous research revealed that genetic loci associated with sitting height ratio show an increased specificity for biological processes related to bone and cartilage, compared to height-associated loci^25^. We observed no significant genetic correlations between ACH-derived facial variation, inflammatory bowel disease^21^ and hormone-sensitive cancer^26^, both serving as “negative control” disorders without known associations to ACH.

**Table 1.** Genetic correlation. Genetic correlation between the ACH-informed GWAS, five ACH-associated traits (body height, infant head circumference, lung volume, obstructive sleep apnea, sitting height ratio), and two negative control traits (hormone-sensitive cancer, cigarettes per day measurement). For each trait, this table contains the Spearman genetic correlation with the ACH-informed GWAS (GC_ACH_), the corresponding 95% confidence interval (CI_ACH_) and pvalue (P_ACH_). We also report the Spearman genetic correlation of these traits with the uninformed facial shape GWAS by White et al. (GC_FACE_), the corresponding 95% confidence interval (CI_FACE_) and p-value (P_FACE_). Genetic correlations that are statistically significant after Bonferroni correction are indicated in bold. Additional information on the selected traits is provided

| Traits | $GC_{ACH}$ | $CI_{ACH}$ | $P_{ACH}$ | $GC_{FACE}$ | $CI_{FACE}$ | $P_{FACE}$ |
| --- | --- | --- | --- | --- | --- | --- |
| **Achondroplasia associated traits** |  |  |  |  |  |  |
| Body height | **0.14** | 0.087-0.20 | 1.2E-07 | **0.23** | 0.17-0.27 | 1.6E-21 |
| Infant head circumference | 0.040 | 0-0.089 | 0.05 | 0.059 | 0.0061-0.11 | 0.01 |
| Lung volume | 0.037 | 0-0.090 | 0.08 | **0.069** | 0.017-0.12 | 0.004 |
| Obstructive sleep apnea | **0.075** | 0.030-0.12 | 5.2E-04 | **0.085** | 0.033-0.14 | 5.9E-04 |
| Sitting height ratio | **0.22** | 0.17-0.27 | 2.6E-21 | **0.24** | 0.20-0.29 | 9.1E-27 |
| **Negative control traits** |  |  |  |  |  |  |
| Hormone-sensitive cancer | 0.024 | 0-0.075 | 0.17 | 0.0030 | 0-0.053 | 0.45 |
| Inflammatory bowel disease | 0.016 | 0-0.067 | 0.27 | 0.007 | 0-0.057 | 0.39 |

### ACH-like phenotype can be obtained in the absence of *FGFR3* mutations

We extended the single SNP analysis to a multivariate genotype-phenotype (MGP) approach that maps the coordinated effects of marker variation of the GWAS associated genes onto craniofacial shape^27,28^. In a sample of 1154 Diversity Outbred mice, the primary MGP associated effect axis resembled an ACH-like phenotype, characterized by a shortened and rounder skull, even when *Fgfr3* marker variation was not included in the model (Fig. 4a-b). Applying the same method to an independent sample of 6772 human cranial vault shapes^29^, the primary MGP associated effect axis revealed a more globular appearance of the cranial vault combined with a relative increase in biparietal diameter (Fig. 4c-d). Similar features are also observed in individuals with ACH, where the narrowing of the skull base can lead to a more rounded calvarium (upper part of the neurocranium). A significant increase in biparietal diameter, but not antero-posterior diameter, of the skull has also been described in individuals with ACH^30^.

## Discussion

Unravelling the complex relationship between genomic and phenotypic variation is a central problem in biology. In this work, we introduce a syndrome-informed phenotyping method to study the connections between the biology of normal-range variation and syndromic variation observed in Mendelian disease, using facial variation in ACH as a case example. Facial features in ACH make up a distinct and recognizable phenotype^15^, represented by the significant facial shape differences between ACH individuals and controls for most parts of the face. In line with existing literature, we found that these differences largely result from changes in phenotypic ‘extremeness’^5,31^. ACH individuals could be well positioned along the extremes of the axes of normal-range facial variation, while some shape deviations remained in those regions of the face that constitute the characteristic ACH facial gestalt. Quantification of ACH-derived facial features in unselected controls showed that individuals vary along the ACH-derived shape axis, and that ACH-like facial shape variation is clearly present in a subset of these control individuals. Though the ACH-derived phenotype axis is derived from a monogenic condition, GWAS of facial shape in an unselected control population revealed a polygenic background for the ACH-derived phenotype scores in this control population. Furthermore, ACH-like craniofacial variation could also be reproduced in two independent datasets of Diversity Outbred mice and control human cranial vaults relying solely on the uncovered polygenic background.

We observed no significant genetic associations with ACH-derived facial shape in the vicinity of the *FGFR3* locus. Similarly, previous GWAS of human facial shape have not found significant associations near *FGFR3*^9^. While the lack of associations between common variants near *FGFR3* and ACH-derived variation does not rule out their contribution, it is also not unexpected given the clinical knowledge on genotype-phenotype correlations for *FGFR3*. Rare variants with large effect in *FGFR3* do not always affect facial features, such as in hypochondroplasia where the mutation causes short stature but no facial dysmorphism^32^. In addition, common variants in *FGFR3* have been associated with idiopathic short stature^33^. This suggests, unlike for height, no general role for *FGFR3* in facial development, but a marked sensitivity to disturbance by very specific large-effect variants. These observations might indicate that facial and skeletal development have different tissue- and/or timepoint-specific sensitivities to disturbances by FGFR3, warranting further research.

Nearly all genes that were identified through the current GWAS had previously been linked to normal-range facial variation^9^, indicating that the ACH-informed phenotype is determined by genes that play a role in facial morphology more broadly. Interestingly, the polygenic background was specifically enriched for biological processes that are disturbed in ACH, such as chondrocyte hypertrophy and differentiation^20^, and genetic correlations were found between ACH-derived facial variation in controls and ACH-linked features such as sitting height ratio^25,34^. These results are in line with previous findings that the effects of major mutations often co-align with the directions of effect linked to broader developmental processes that are affected by those mutations^27^.

The convergence of genetic effects onto shared axes of shape variation stems from the highly integrated nature of the human face^35,36^. While myriad genes can influence facial morphology, the potential directions (axes) in which facial shape can vary are constrained by the developmental processes on which they act^35^. Our findings indicate that both normal human facial variation and facial variation associated with rare Mendelian syndromes occur along the same developmental axes. These developmental axes appear to be determined by a background of common polygenic variation. Rare Mendelian genetic variants with major effects appear to move individuals further toward the extreme end of these axes. The polygenic variation that underlies these developmental axes thus likely contributes to the range of variation seen in the corresponding Mendelian syndromes. Indeed, this may partially explain the occurrence of subclinical phenotypes in conditions such as orofacial clefting^37^, as well as a tendency for unaffected relatives of probands with craniofacial syndromes to sometimes themselves be misclassified as syndromic by an automated syndrome classification tool based on 3D facial imaging^38^.

The finding that normal and syndromic facial variation are related through shared developmental axes also has implications for the mechanisms of variable expressivity and penetrance. This phenomenon likely occurs because developmental processes drive directions of variation on which multiple genomic and environmental influences may converge. For ACH, this would mean that the *FGFR3* gain-of-function mutation produces a large-scale effect on an axis of variation that exists in the general population, and is driven by variation in growth at the cranial synchondroses and cartilaginous growth centers in early craniofacial development. Mutations that alter chondrocyte proliferation or maturation in mice show directions of effect that broadly resemble ACH including doming of the neurocranium, decreased cranial base flexion and reduction in midfacial prognathism^39,40^. If individuals with ACH vary along this same multivariate axis of facial shape, modulation of the degree of cartilage proliferation could explain variation in phenotypic severity for ACH. Conversely, when individuals with ACH vary in directions orthogonal to this axis, however, this would point towards other developmental drivers of variation^35^.

The value of integrating common and Mendelian disease genetics was recently demonstrated by Blair et al.^5^, who mapped heterogeneous symptom data to latent quantitative traits for various Mendelian diseases. Genomic association testing of the newly derived traits revealed common variants predictive of disease outcome; however, the inference of latent traits required phenotypes available at biobank-scale, limiting applicability of that approach. The syndrome-informed framework we present here is applicable to many other phenotypes and, importantly, to relatively small sample sizes, which remains a major challenge in studies of rare diseases. Here, we studied ACH (n = 43) as proof of principle, but the syndrome-informed framework can also be generalized to other genetic disorders. For example, genomic analysis of Pierre Robin Sequence-derived phenotypic scores identified genetic variants near the *SOX9* locus, which is commonly linked to the disorder, among other genetic loci that are thought to conjointly modulate the facial phenotype^41^. In addition, applying our approach to genetic conditions with a poorly understood pathophysiology could highlight developmental and biological pathways of importance. Similarly, by defining a shape axis based on a group of individuals with similar phenotypic features but unknown diagnosis, our approach could provide insight into the shared genetic etiology and impaired pathways in these individuals. And, of particular importance, the polygenic background identified using a syndrome-informed approach may highlight interesting targets to identify putative modifiers of phenotypic expression in monogenic disorders^42^.

In conclusion, genetically mapping ACH-derived phenotypic effects in the general population highlighted a polygenic basis for a shape axis determined by a monogenic disorder. Jointly modeling these candidate genes in turn revealed that ACH-like phenotypes can be generated without *FGFR3*. These findings have important implications for unravelling the relationship between discrete and continuous variation and for understanding the role of causative genes for Mendelian disorders. If causative genes act on already existing axes of variation determined by developmental processes, then they are causes in only a limited sense in the background developmental context. Disease associated variants may be more productively seen as belonging to a larger set of potential perturbations capable of shifting phenotypes along developmentally determined directions of variation. This framework also promotes understanding of variable expressivity and penetrance in genetic disease, which is of great value to aid diagnosis and improve patient outcomes.

## Methods

### Sample composition

We obtained 3D facial photos, demographic data (age, sex, self-reported ethnicity) and clinical and/or molecular testing results of 70 individuals with achondroplasia from the online FaceBase repository (www.facebase.org; FB00000861). From this group we excluded individuals of self-reported non-European descent (n = 21) and those with incomplete or missing metadata (n = 6) to retain a curated sample of 43 individuals.

The unselected control sample consisted of 3D facial images, demographics (age, sex, genomic ancestry) and imputed genotype data of 8246 unrelated individuals of European descent originating from the United States (US) and the United Kingdom (UK)^19^. The US dataset (n = 4680) included samples from the 3D Facial Norms cohort^43^ and studies at the Pennsylvania State University and Indiana University-Purdue University Indianapolis. The UK sample (n = 3566) consisted of participants from the Avon Longitudinal Study of Parents and Children (ALSPAC)^44,45^. European participants were identified by projecting them into a principal component space constructed using the 1000G Phase 3 dataset. Participants with missing covariate information (e.g., age, sex) or with insufficient image quality were excluded. Detailed sample characteristics and further information on the calculation of the genetic ancestry axes are provided by White et al.^19^.

The appropriate local ethical approvals were obtained, and all participants gave written informed consent prior to participation. Ethical approval for the ALSPAC study was obtained from the ALSPAC Ethics and Law Committee and the Local Research Ethics Committees.

### Genotyping and imputation

Genotyping and imputation of the European control sample was performed as described previously^19^. In brief, genotypes of the three different US subsamples, separately, were phased using SHAPEIT2 (v2.r900)^46^ and imputed to the 1000 Genomes Phase 3 reference panel^47^ using the Positional Burrows-Wheeler Transform pipeline (v3.1)^48^ of the Sanger Imputation Server (v0.0.6)^49^. SNP-level (INFO score < 0.8) and genotype per participant-level (genotype probability < 0.9) filters were used to omit poorly imputed variants. Finally, a single US cohort was obtained by merging the subsamples and filtering the SNPs based on missingness across individuals (--geno 0.5), minor allele frequency (--maf 0.01), and Hardy-Weinberg equilibrium (p < 1e-6), resulting in 7,417,619 SNPs for analysis.

For the UK dataset, imputed genotypes were obtained directly from the ALSPAC database. SHAPEIT2^46^ was used for pre-phasing of haplotypes and imputation against the 1000 Genomes Phase 1 reference panel (Version 3)^50^ was performed using IMPUTE2^51^. After post-imputation quality control, the UK dataset contained 8,629,873 SNPs for analysis. Because restrictions are in place against merging the ALSPAC genotypes with any other genotypes, these were held separately during the analysis.

In total, 7,417,619 SNPs were overlapping between US and UK datasets, which were used in subsequent genetic association analyses. SNPs on the X chromosome were coded 0/2 for hemizygous males, to match with the 0/1/2 coding for females.

### Facial phenotyping and segmentation

3D facial images were acquired using three digital stereophotogrammetry systems (3dMDface, Vectra H1, Creaform Gemini) and one laser scanning system (Konica Minolta VI-900). We non-rigidly registered an average facial atlas to each 3D image using the MeshMonk toolbox^16^ to obtain a standard facial representation defined by 7160 homologous quasi-landmarks. Due to the bilaterally paired construction of the quasi-landmarks constituting the atlas, registered images were symmetrized by averaging the original configuration and its horizontally reflected copy following Procrustes superimposition. Images were visually inspected and excluded if the registration process had failed.

All symmetrized quasi-landmark configurations of the unselected control sample were aligned by generalized Procrustes analysis (GPA) and adjusted for sex, age, and age-squared in a partial least-squares regression (PLSR, function plsregress in MATLAB). Facial shape was divided into 63 global-to-local segments by hierarchical spectral clustering as described elsewhere^17^, providing facial segments at five hierarchical levels of scale. In each segment separately, symmetrized and covariate-adjusted shapes were aligned using GPA and dimensionality was reduced by principal component analysis (PCA), with the optimal number of principal components (PC) to retain determined by parallel analysis. We normalized the projections on each PC to have unit variance by dividing each projection by the standard deviation of all projections.

Next, the ACH sample was superimposed onto the mean control shape using GPA and shapes were corrected for the same sex and age covariates using the regression coefficients from the PLSR model of the control sample. We then applied the same facial segmentation to the ACH sample and participants were projected into each segment-specific principal component space, again normalizing by dividing each projection by the standard deviation of all projections from the control sample. To assess how much of the facial variation was retained by projecting ACH syndromic samples into a principal component space derived from unselected, non-syndromic individuals, we measured the reconstruction error between the age-and sex-corrected shapes and their corresponding projections as the mean Euclidean distance across all 7160 quasi-landmarks.

### Achondroplasia-informed phenotyping

For each of the 63 facial segments separately, an ACH-derived facial trait was defined as follows. First, in the variation standardized space, we established an ACH facial trait as the shape axis passing through the average PC projection of the control sample and the ACH average projection. We then obtained univariate trait scores for each control subject by computing the cosine of the angle between their individual vector, going from the average PC projection of the control sample to their individual PC projections, and the ACH trait vector^52,53^. These scores were computed in a leave-one-out scheme such that each subject was excluded from learning the trait vectors on which they were scored. To evaluate the ACH-like effect size in the control population, we regressed global facial shape onto the trait scores obtained for segment 1 (full face) using PLSR and report R-squared as the percentage of phenotypic variation explained.

We additionally tested whether ACH facial shapes differed significantly from a matched control sample of equal size. In a random order, we matched each ACH sample to a control sample of the same sex that was closest in age. The selected control was then omitted from the pool of potential matches. We co-aligned the covariate-adjusted and symmetrized quasi-landmarks of both groups using GPA and regressed facial shape onto group membership using PLSR. A p-value was generated by a permutation test on R-squared with 10,000 permutations. This was done for each segment separately, and significant differences (p < 0.05) were observed in 58 out of 63 facial segments.

### Genome-wide association study

For both US and UK datasets separately, we combined the ACH-derived trait scores across the 58 significant segments into a single phenotype matrix ([n x m] with n_US_ = 4680 controls, n_UK_ = 3566 controls, and m = 58 facial segments). This phenotype matrix was tested for genome-wide SNP-associations in a multivariate association framework using canonical correlation analysis (CCA) following White et al.^16^. However, instead of performing a separate GWAS per facial segment, scores generated across multiple segments were now combined into a single multivariate GWAS. The GWAS was conducted following a two-stage design with both US and UK cohorts alternating as the discovery and replication sets. First, we applied CCA in the discovery sample to obtain association p-values as well as the shape axis maximally correlated with each SNP. Next, the replication sample was projected onto this axis to ensure consistency of the phenotype, leading to univariate trait scores which were subsequently tested for genetic associations in a linear regression model. Finally, discovery and replication p-values were aggregated in a meta-analysis using Stouffer’s method^54^. Per SNP, the GWAS design generated two meta-analysis p-values, meta_US_ and meta_UK_, reflecting the sample that served as the discovery set. Because CCA does not accommodate adjustments for covariates, we corrected the dependent (facial shape) and independent variables (genotypes) for age, age-squared, sex, height, weight, facial size, four genomic ancestry axes, and camera system using PLSR prior to GWAS.

The lowest meta-analysis p-value per SNP was selected and compared against the genome-wide Bonferroni threshold (p < 5e-8). We observed 1925 SNPs at the level of genome-wide significance, which were clumped into 35 independent loci as follows. Starting from the lead SNP (lowest p-value), SNPs within 10kb or within 1Mb but in linkage disequilibrium (r^2^ > 0.01) were clumped into the same locus represented by the lead SNP. Next, considering the lead SNPs only, signals within 10Mb and r^2^ > 0.01 were merged. Third, any locus with a singleton lead SNP was removed. For each of the lead SNPs, the nearest gene was assigned as the candidate gene.

In the multivariate GWAS setup, CCA extracts the linear combination of the ACH trait scores for the 58 significant facial segments that maximally correlates with the SNP being tested. From the CCA loadings, we examined which of the facial segments contributed most to the observed GWAS signals to delineate the associated shape effects.

### Protein network analysis

We searched the STRING database^55^ for known interactions with FGFR3. We focused on high-confidence interactions (confidence score 0.7) derived from curated databases or experimentally determined (Supplementary Table 2). SNP p-value data were aggregated to gene-association scores (gene-level p-values) and we evaluated enrichment for associations with the FGFR3 network using MAGMA^56^. Next, we performed protein-protein interaction analysis of the GWAS candidate genes and evaluated potential associations, direct or indirect, with the FGFR3 network using default settings (confidence score 0.4, including all interaction sources).

### Gene set enrichment analysis

We used GREAT^18^ to associate the 35 genetic loci to Gene Ontology (GO) annotations and calculated enrichment of biological processes for these annotations. To assess targeted enrichment of processes specific to our ACH-informed phenotyping approach, we compared gene set enrichment of all biological processes that reached significance in the hypergeometric test to a recent GWAS of normal facial variation in the current European control sample by White et al.^19^. In addition, we repeated GO term enrichment against three background sets of craniofacial-associated genes as summarized by Naqvi et al.^9^. The first set consisted of all genes identified in 25 previously published GWASs of facial shape, the second set contained genes with known roles in Mendelian craniofacial disorders and/or orofacial clefting, and the third set is a combination of both GWAS and disease-associated genes. As a negative control, we repeated the analysis with results from a recent GWAS of inflammatory bowel disease as foreground set^21^.

A database of genes and annotated ontology terms was downloaded from the StringDB website (https://stringdb-static.org/download/protein.enrichment.terms.v11.5.txt.gz). For each term, *τ*, a hypergeometric p-value was calculated as

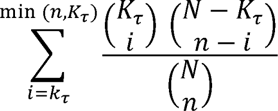

with *N* the total number of genes in the background set, *n* the total number of genes in the foreground set, *K* the number of genes with annotation *τ* in the background set, and *k* the number of genes with annotation *τ* in the foreground set. P-values were adjusted to a 5% false discovery rate (FDR) using the Benjamini-Hochberg procedure^57^.

### Gene expression in chondrocytes

We computed gene-level p-values (gene-association scores) based on the ACH-informed GWAS summary statistics using MAGMA^56^, as well as from published GWAS summary statistics of normal facial shape in the same unselected control population^19^, height^58^, and inflammatory bowel disease^21^.

We downloaded published microarray data from murine growth plate dissections from the GEO data repository^59^ (accession number GSE87605). Probe identifiers were mapped to mouse genes using the Mouse Genome Informatics database^60^. Mouse gene names were subsequently mapped to their human homologs using the Ensembl BioMaRt tool^61^. We calculated gene expression specificity scores per epiphyseal layer by dividing the expression of each gene per layer by the total expression of that gene. We calculated gene expression Z-scores per chondrocyte maturation stage by averaging gene expression across the four available samples for day 3 of embryonic development (early maturation stage) and day 10 of embryonic development (late maturation stage). Finally, we evaluated the Pearson correlation between gene-level p-values, specificity scores for expression per epiphyseal layer, and Z-scores per chondrocyte maturation stage.

From the GEO data repository^59^, we downloaded murine chondrocyte RNAseq data (accession number GSE225796) and used the DESeq function (DESeq2 package) in R to perform differential gene expression analysis. DESeq2 transforms read counts based on size factors and dispersion, fits a negative binomial generalized linear model (GLM), performs a Wald significance test, and assesses differentially expressed genes based on a false discovery rate cutoff of 0.05 using the Benjamini-Hochberg procedure.

### Genetic correlation

To assess the extent to which genome-wide profiles of association were shared with known ACH related traits, we computed the Spearman correlation between two vectors of linkage disequilibrium (LD)-block stratified association p-values. This approach provides a multivariate, robust alternative to LD score regression (LDSC)^62,63^ for computing genetic correlations and is applicable to unsigned summary statistics yielded by CCA^22^. We collected publicly available genome-wide summary statistics for five traits with known associations with the achondroplasia phenotype (body height^58^, head circumference^64^, lung volume^65^, obstructive sleep apnea syndrome^66^ and sitting height ratio^67^) and for two putative unrelated traits (hormone-sensitive cancer^26^ and inflammatory bowel disease^21^) to serve as negative controls. Details on the selected traits and links to relevant publications are summarized in Supplementary Table 5. LD scores were readily obtained from the 1000 Genomes European data^46^ and SNPs were filtered to HapMap3 SNPs, excluding SNPs in the Major Histocompatibility Complex region^68^. For each LD block, we computed the mean SNP −log10(p-value) and computed a rank-based Spearman correlation using the average association value for that LD-block. We estimated the standard error of the Spearman correlation using a bootstrapping approach with 100 resampling cycles.

### Multivariate genotype-phenotype mapping

We applied the GWAS candidate genes to the recent multivariate genotype-phenotype (MGP) model in Diversity Outbred (DO) mice^27^ in R (version 4.2.0). Composition (n = 1154 samples), genotyping (n = 123,309 markers) and landmarking (n = 54 3D landmarks) of the DO sample is described in detail by Aponte et al.^27^. In a regularized partial least squares model, the MGP method identifies axes of shape variation that maximally covary with genetic marker variation for the selected gene set. The regularization parameter was determined at 0.06 based on 10-fold cross-validation. For each of the genes, the MGP model outputs their overall contribution, or marker loadings, to the estimated shape axes. The principal axis of shape covariation is visualized directly onto the mouse craniofacial shape as a heatmap, representing the displacement along the surface normals with reference to the mean DO shape.

For 6,772 multi-ancestry participants of the Adolescent Brain Cognitive Development (ABCD) study^69^, the outer head surface was extracted from magnetic resonance images as described by Goovaerts et al.^29^. The MeshMonk toolbox^16^ was used to perform rigid and subsequently nonrigid surface registration using a full-head template comprising 28,218 quasi landmarks. From this, we cropped out the area covering the cranial vault (n = 11,410 quasi landmarks), encompassing the supraorbital ridge, and extending towards the occipital bone. Shapes were then adjusted for age, sex, weight, height, cranial size, scanner site, and the first 10 genomic PCs using PLSR after GPA alignment. Following PCA and parallel analysis, 65 orthogonal axes of cranial vault shape variation were retained and normalized to unit variance. Using CCA, we optimized the linear combinations of ACH lead SNPs and vault shape PCs to extract a maximally correlated latent phenotype. For SNPs not found in the ABCD sample, we searched for proxy SNPs within the 1000 Genomes Phase 3 European sample and selected the SNP in strongest LD with the original SNP and with at least r^2^ > 0.9. No suitable proxy was available for rs113434679. The latent phenotypic traits were visualized directly onto the head surface as a heatmap, representing the displacement along the surface normals with reference to the mean head surface.

## Ethics statement

This study was approved by the ethical review board of KU Leuven and University Hospital Leuven (S60568, Leuven, Belgium), and the University of Calgary (REB14-0340, Calgary, Canada). Local institutional approval was granted for access to the FaceBase Repository (S60658, Leuven, Belgium). The work on mouse craniofacial shape was performed according to protocols approved and reviewed by animal care committees at the University of Calgary (AC13-0268) and the University of Alberta (AUP1149).

## Funding

This work was supported by the Research Fund KU Leuven (BOF-C1, C14/15/081 & C14/20/081 to PC); the Research Program of the Research Foundation-Flanders (FWO, G0D1923N to PC); NIH-NIDCR (R01-DE027023 to SMW; U01DE024440 to RAS, ODK and BH)

### Achondroplasia sample

FaceBase data collection and analyses were supported by NIH-NIDCR (U01DE024440 to RAS, ODK and BH).

### Unselected control sample

Pittsburgh personnel, data collection and analyses were supported by the National Institute of Dental and Craniofacial Research (U01-DE020078 to MLM and SMW; R01-DE016148 to MLM and SMW; R01-DE027023 to SMW). Funding for genotyping by the National Human Genome Research Institute (X01-HG007821 & X01-HG007485 to MLM).

Penn State personnel, data collection and analyses were supported by the Center for Human Evolution and Development at Penn State, the Science Foundation of Ireland Walton Fellowship (04.W4/B643 to MDS), the US National Institute of Justice (2008-DN-BX-K125 to MDS; 2018-DU-BX-0219 to SW) and by the US Department of Defense.

IUPUI personnel, data collection and analyses were supported by the National Institute of Justice (2015-R2-CX-0023, 2014-DN-BX-K031 & 2018-DU-BX-0219 to SW).

The UK Medical Research Council and Wellcome (grant no. 102215/2/13/2) and the University of Bristol provide core support for ALSPAC. The publication is the work of the authors and they will serve as guarantors for the contents of this paper. A comprehensive list of grants funding is available on the ALSPAC website (http://www.bristol.ac.uk/alspac/external/documents/grant-acknowledgements.pdf). ALSPAC GWAS data was generated by Sample Logistics and Genotyping Facilities at Wellcome Sanger Institute and LabCorp (Laboratory Corporation of America) using support from 23andMe. We are extremely grateful to all the families who took part in this study, the midwives for their help in recruiting them, and the whole ALSPAC team, which includes interviewers, computer and laboratory technicians, clerical workers, research scientists, volunteers, managers, receptionists and nurses.

### Human cranial vault

Data on human cranial vault shape were obtained from the Adolescent Brain Cognitive Development (ABCD) Study (https://abcdstudy.org), held in the NIMH Data Archive (NDA). This is a multisite, longitudinal study designed to recruit more than 10,000 children aged 9-10 and follow them over 10 years into early adulthood. The ABCD Study is supported by the National Institutes of Health and additional federal partners under award numbers U01DA041048, U01DA050989, U01DA051016, U01DA041022, U01DA051018, U01DA051037, U01DA050987, U01DA041174, U01DA041106, U01DA041117, U01DA041028, U01DA041134, U01DA050988, U01DA051039, U01DA041156, U01DA041025, U01DA041120, U01DA051038, U01DA041148, U01DA041093, U01DA041089, U24DA041123, U24DA041147. A full list of supporters is available at https://abcdstudy.org/federal-partners.html. A listing of participating sites and a complete listing of the study investigators can be found at https://abcdstudy.org/consortium_members/. ABCD consortium investigators designed and implemented the study and/or provided data but did not necessarily participate in analysis or writing of this report. This manuscript reflects the views of the authors and may not reflect the opinions or views of the NIH or ABCD consortium investigators.

The ABCD data repository grows and changes over time. The ABCD data used in this report came from 10.15154/1519007. DOIs can be found at http://dx.doi.org/10.15154/1519007.

### Sitting height ratio GWAS

We thank Eric Bartell, Joanne Cole, and Joel Hirschhorn for their contribution of pre-publication GWAS of sitting height ratio summary statistics to this work.

---

**Supplementary Figure 1.**
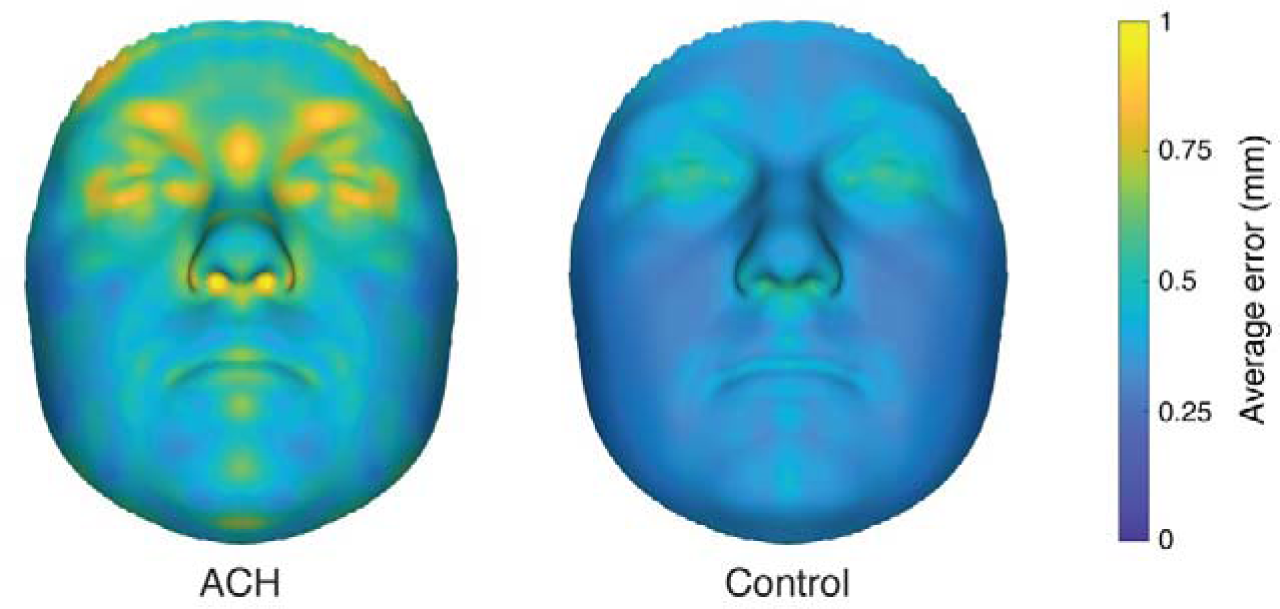
Average reconstruction error. Reconstruction error (in mm) between the original quasi-landmark configurations and their corresponding projections in principal component space (constructed from healthy controls only), averaged across all ACH and control samples, respectively.

**Supplementary Figure 2.**
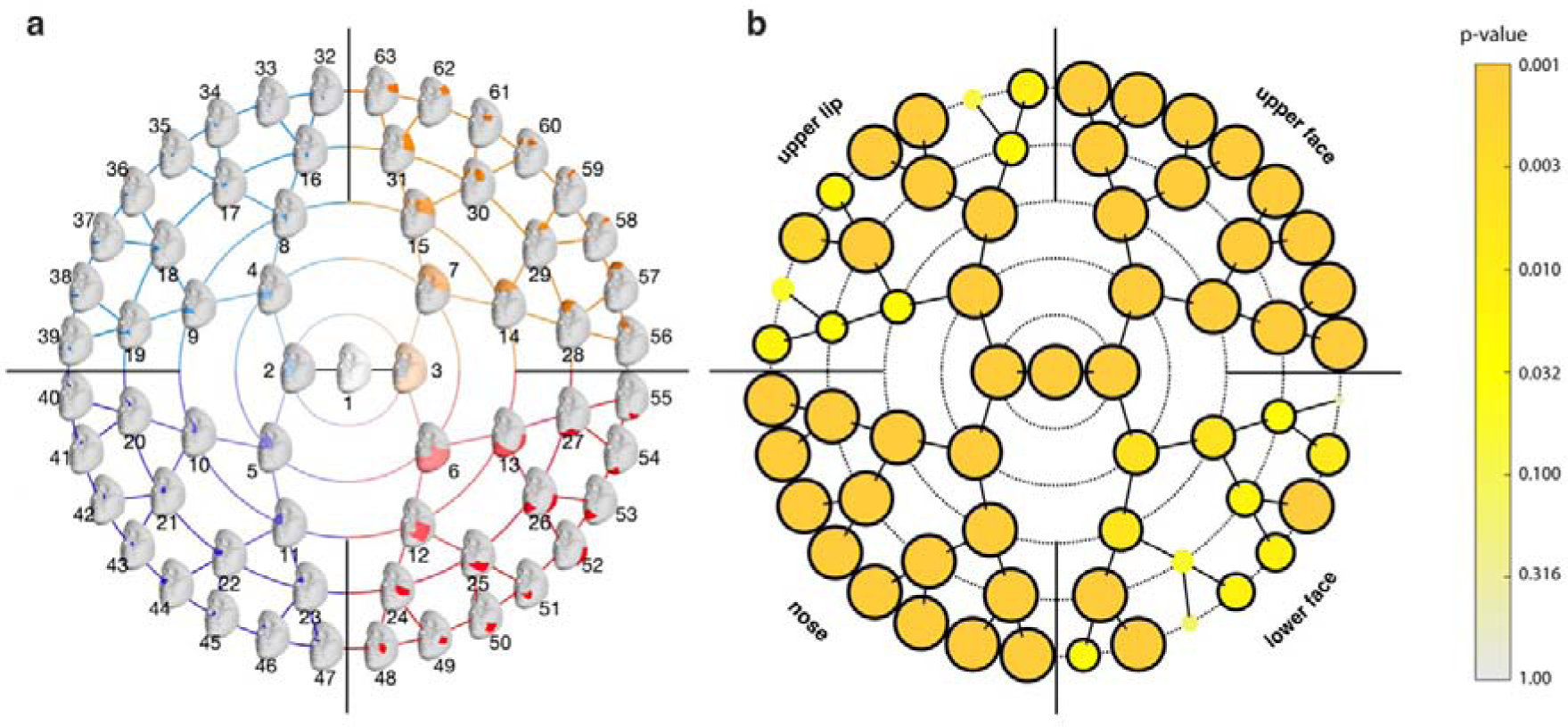
Global-to-local segmentation. **(a)** 63 facial segments obtained by hierarchical spectral clustering, ordered and colored according to facial quadrant. Starting from the full face in the center, each segment is subdivided into two smaller segments, providing a detailed description of facial shape at multiple levels of scale. **(b)** For each of the 63 segments, we tested if facial shape was significantly different between the ACH and control samples in an age- and sex-matched setup. Nodes are colored according to the p-value, with black outlines indicating statistical significance (p < 0.05). The position of each node corresponds to the facial segments depicted in panel a.

**Supplementary Figure 3.**
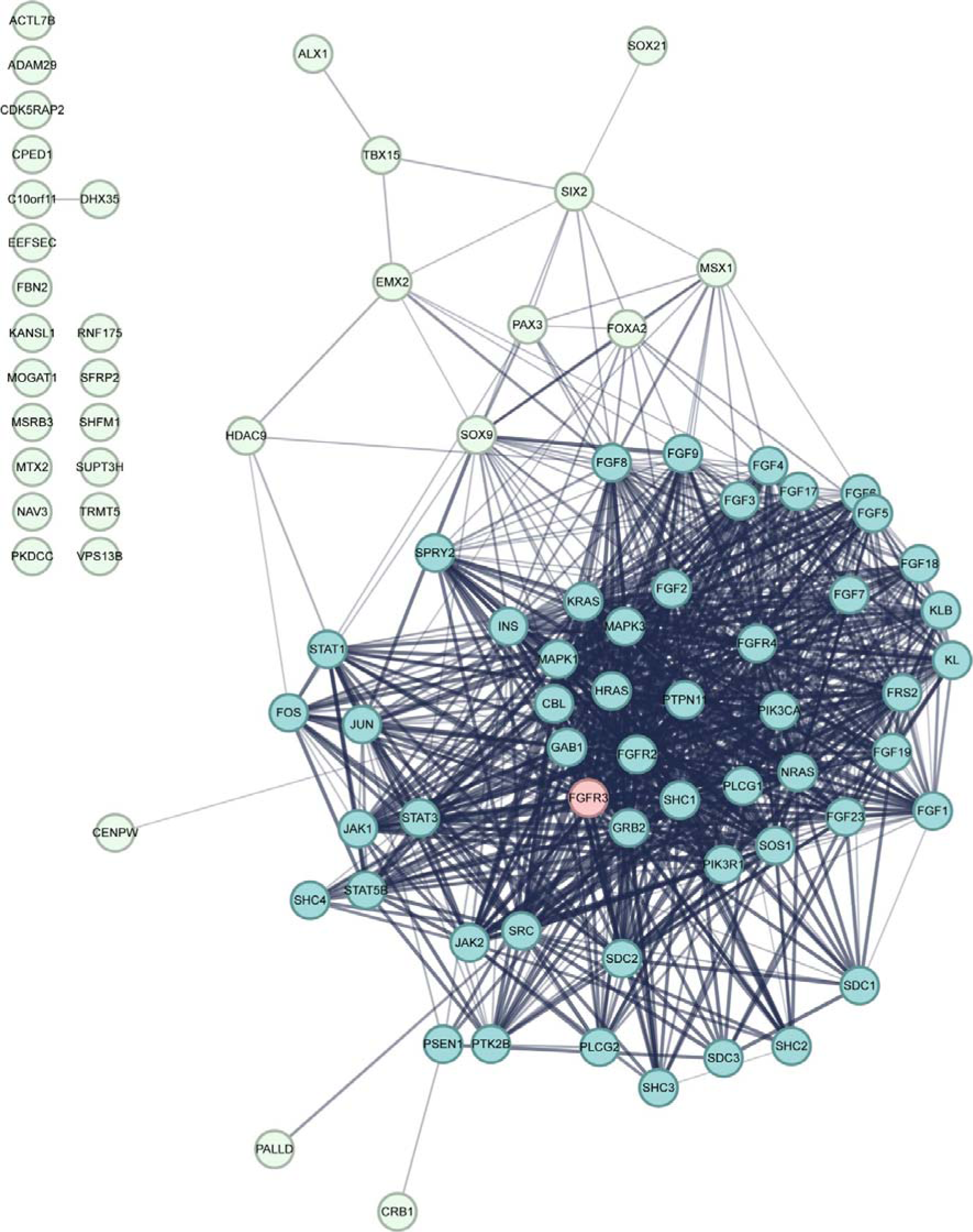
Predicted FGFR3 interaction network. High confidence interactions (confidence score > 0.7) are shown in blue; the candidate genes identified through GWAS are shown in green. Line thickness indicates the degree of confidence prediction of the interaction.

**Supplementary Figure 4.**
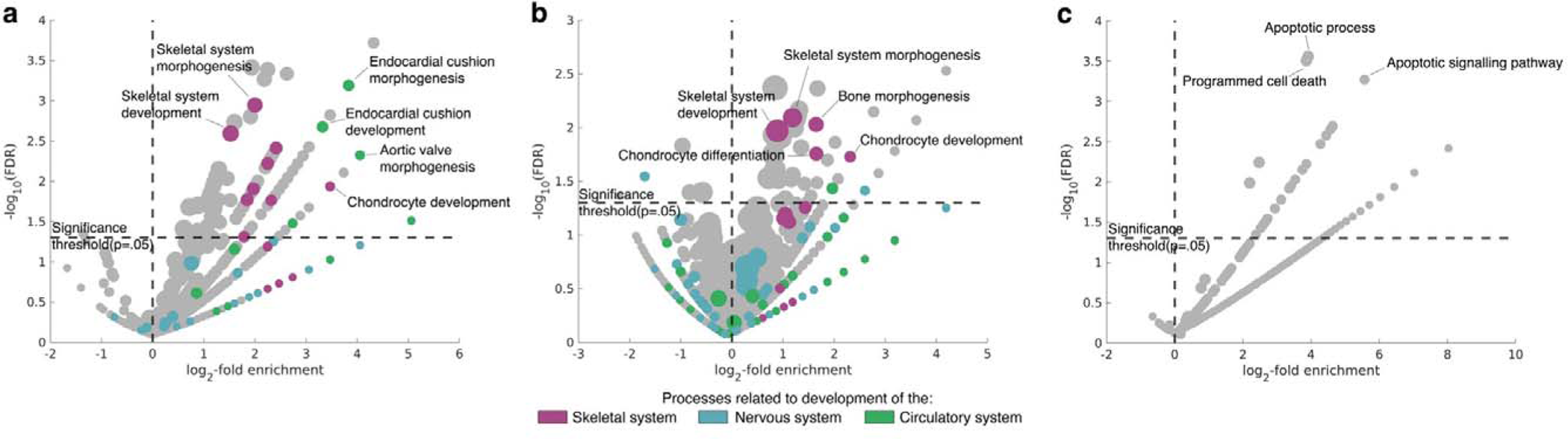
GO enrichment analysis. Fold enrichment of Gene Ontology (GO) biological processes enriched in the ACH GWAS compared to different background sets. **(a)** ACH-informed GWAS versus genes implicated in Mendelian craniofacial disorders **(b)** ACH-informed GWAS versus all genes previously identified through GWAS of facial shape and genes implicated in Mendelian craniofacial disorders combined **(c)** Negative control GWAS of inflammatory bowel disease versus all genes previously identified through GWAS of facial shape and genes implicated in Mendelian craniofacial disorders. Only processes enriched in both studies are displayed. Node size corresponds to the number of genes mapped to each process.

