## Supplementary Data 1 for "Syndrome-informed phenotyping identifies a polygenic background for achondroplasia-like facial variation in the general population"

**Supplementary Data 1. Principal component score distributions.** PC scores of the ACH (in green) and control (in beige) samples on the first 65 PCs (determined by parallel analysis). The PCA space was constructed from the control sample, and ACH samples were then projected into this space. The facial shape changes associated with each PC are depicted by the facial morphs, scaled to 3 times the standard deviation of the PC scores. The proportion of variance explained by each PC is listed between brackets.

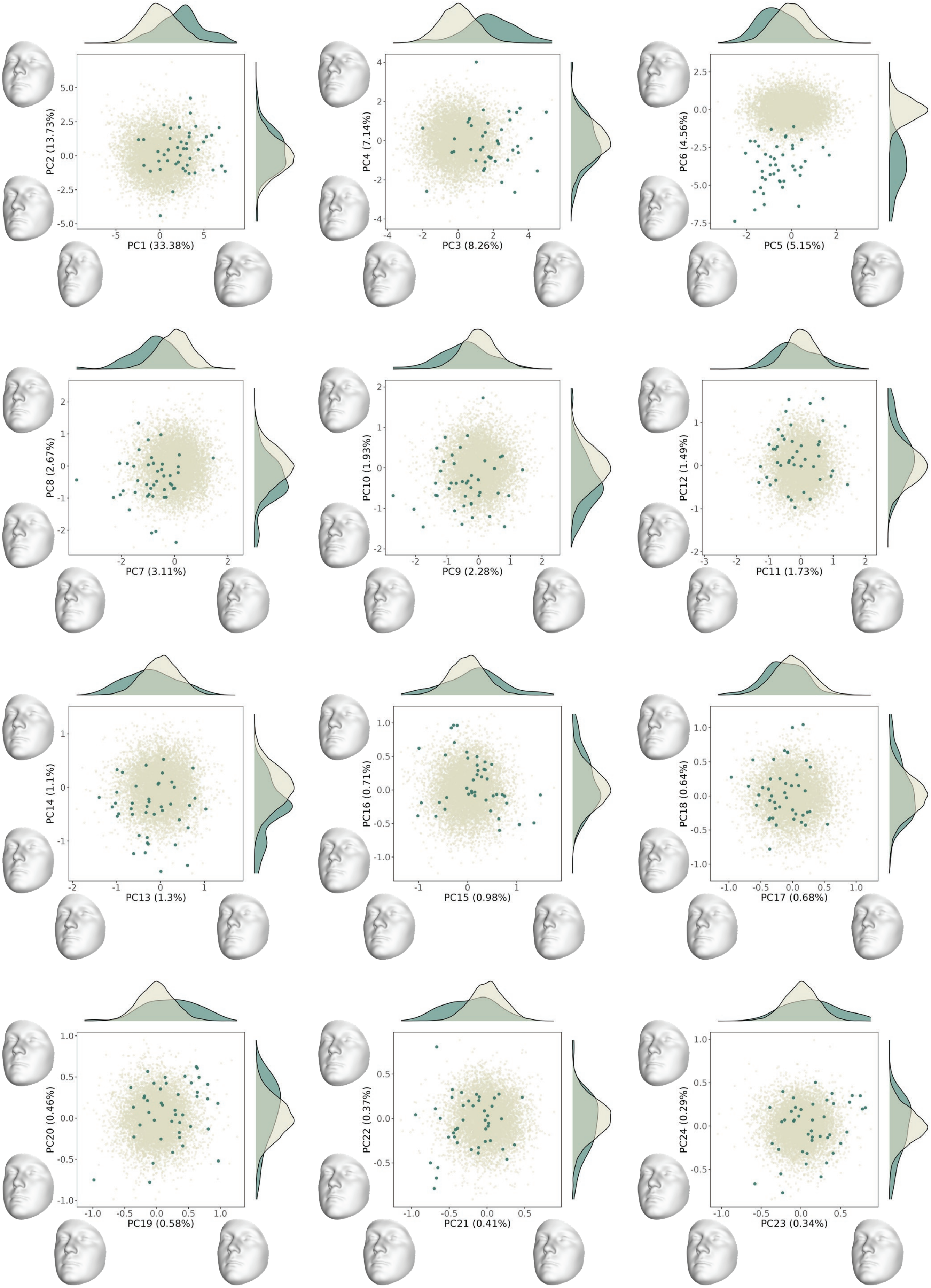

PC1 (33.38%)
PC2 (13.73%)
PC3 (8.26%)
PC4 (7.14%)
PC5 (5.15%)
PC6 (4.56%)
PC7 (3.11%)
PC8 (2.67%)
PC9 (2.28%)
PC10 (1.93%)
PC11 (1.73%)
PC12 (1.49%)
PC13 (1.3%)
PC14 (1.1%)
PC15 (0.98%)
PC16 (0.71%)
PC17 (0.68%)
PC18 (0.64%)
PC19 (0.58%)
PC20 (0.46%)
PC21 (0.41%)
PC22 (0.37%)
PC23 (0.34%)
PC24 (0.29%)

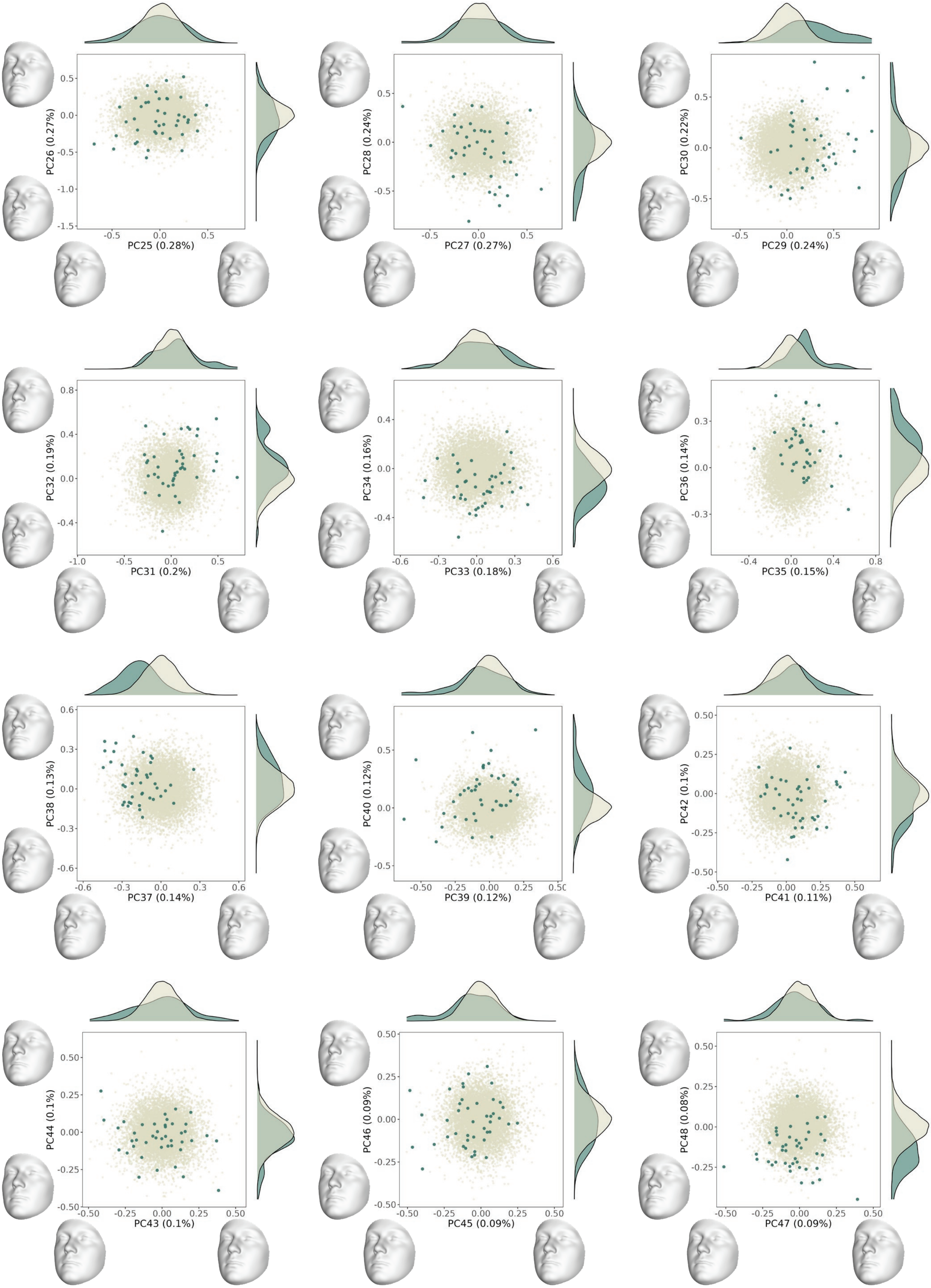
PC25 (0.28%)
PC26 (0.27%)
PC27 (0.27%)
PC28 (0.24%)
PC29 (0.24%)
PC30 (0.22%)
PC31 (0.2%)
PC32 (0.19%)
PC33 (0.18%)
PC34 (0.16%)
PC35 (0.15%)
PC36 (0.14%)
PC37 (0.14%)
PC38 (0.13%)
PC39 (0.12%)
PC40 (0.12%)
PC41 (0.11%)
PC42 (0.1%)
PC43 (0.1%)
PC44 (0.1%)
PC45 (0.09%)
PC46 (0.09%)
PC47 (0.09%)
PC48 (0.08%)

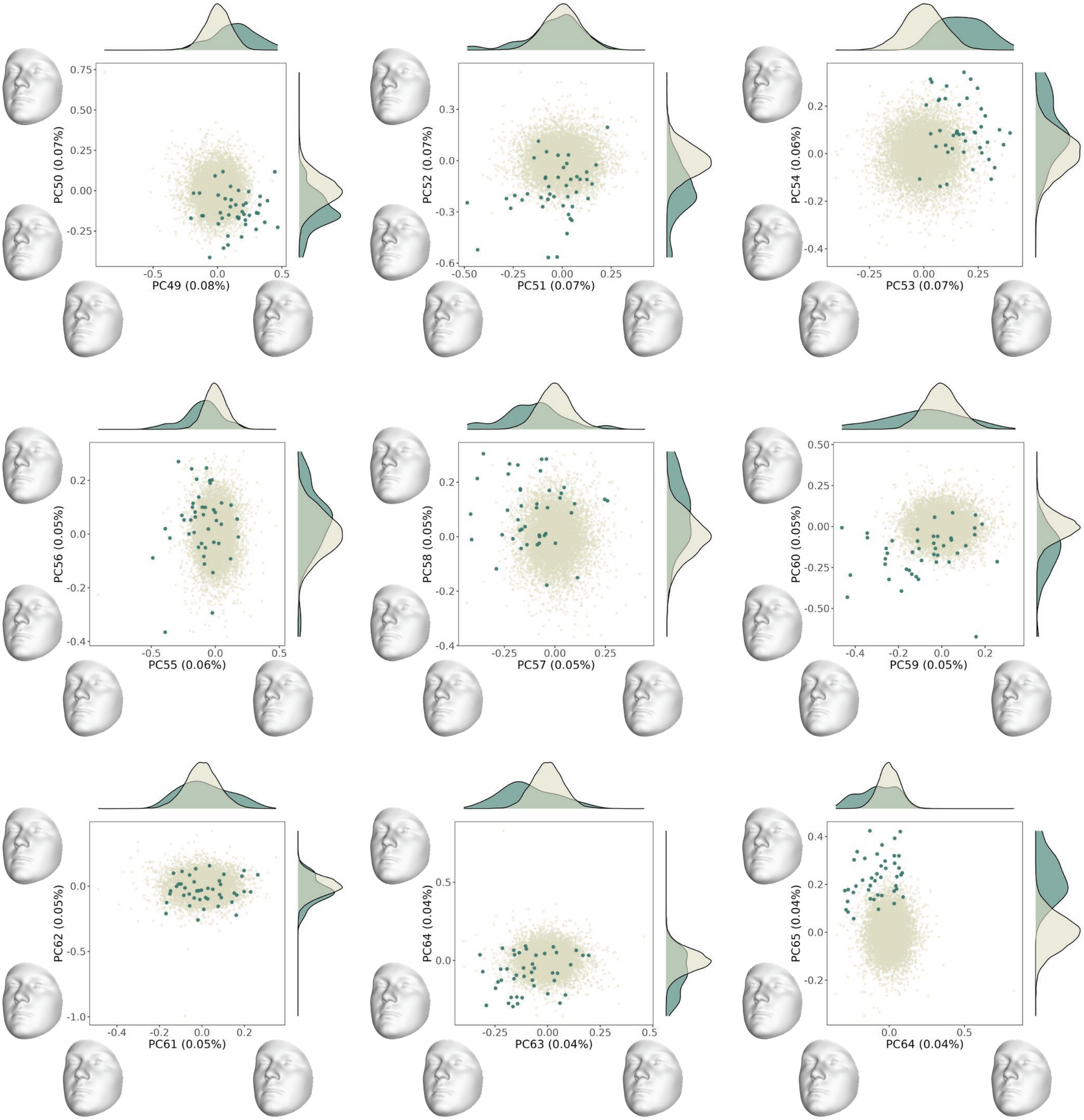

0.75
0.50
0.25
0.00
-0.25
-0.5
0.0
0.5
PC50 (0.07%)
PC49 (0.08%)
0.3
0.0
-0.3
-0.6
-0.50
-0.25
0.00
0.25
PC52 (0.07%)
PC51 (0.07%)
0.2
0.0
-0.2
-0.4
-0.25
0.00
0.25
PC54 (0.06%)
PC53 (0.07%)
0.2
0.0
-0.2
-0.4
-0.5
0.0
0.5
PC56 (0.05%)
PC55 (0.06%)
0.2
0.0
-0.2
-0.4
-0.25
0.00
0.25
PC58 (0.05%)
PC57 (0.05%)
0.50
0.25
0.00
-0.25
-0.50
-0.4
-0.2
0.0
0.2
PC60 (0.05%)
PC59 (0.05%)
0.0
-0.5
-1.0
-0.4
-0.2
0.0
0.2
PC62 (0.05%)
PC61 (0.05%)
0.5
0.0
-0.25
0.00
0.25
0.5
PC64 (0.04%)
PC63 (0.04%)
0.4
0.2
0.0
-0.2
0.0
0.5
PC65 (0.04%)
PC64 (0.04%)
