## Supplementary Data 2 for "Syndrome-informed phenotyping identifies a polygenic background for achondroplasia-like facial variation in the general population"

**Supplementary Data 2. Trait score distributions per facial segment.** Probability density functions of the ACH trait scores (individual PC projection scores on the ACH trait vector) per facial segment, plotted for both control (in beige) and ACH datasets (in green). Values smaller than 1 indicate more ACH-like; values greater than 1 indicate less ACH-like. The facial morphs on the bottom depict the shape effects associated with the ACH-derived trait axes per segment.

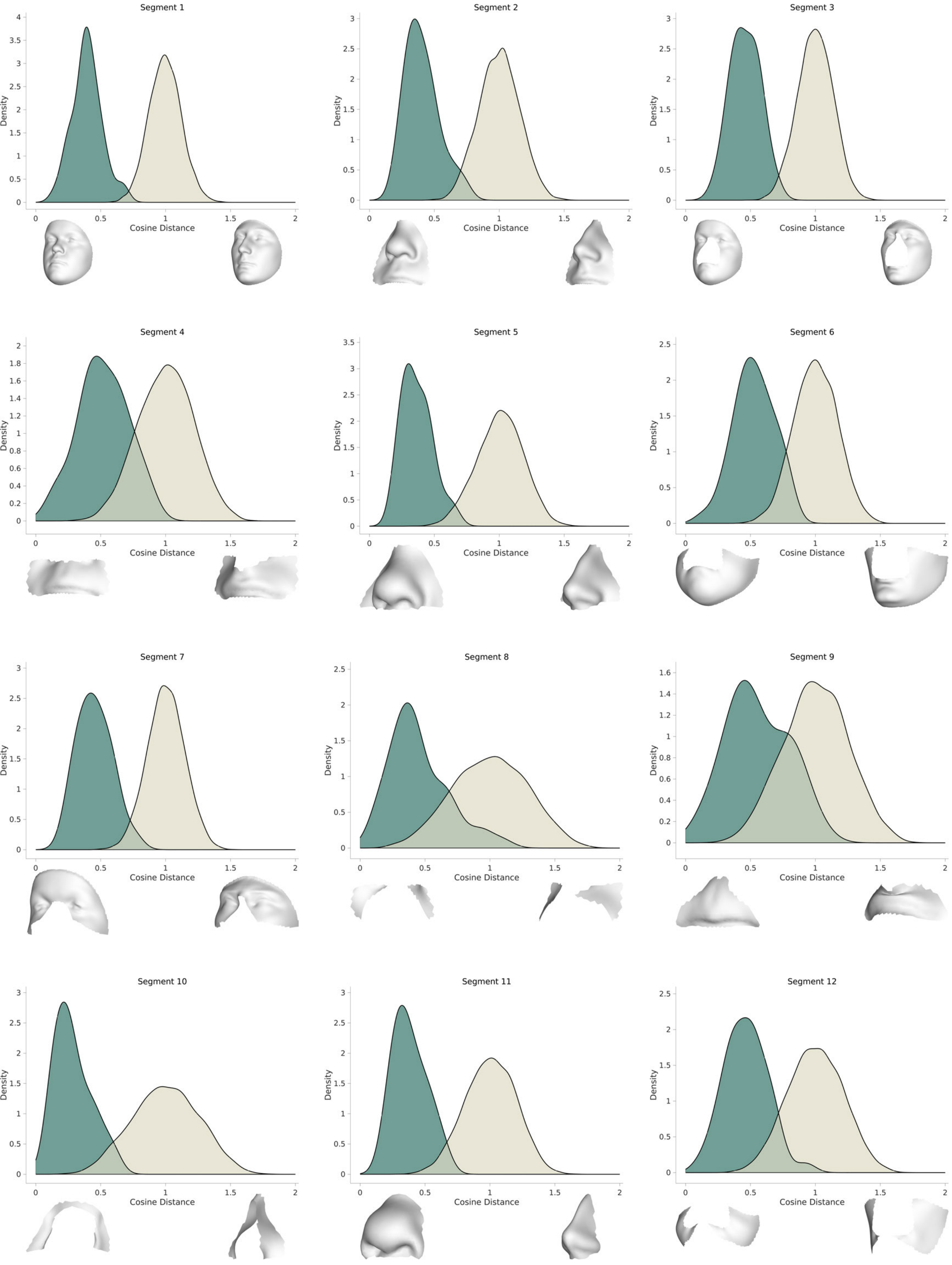
Segment 1
Segment 2
Segment 3
Segment 4
Segment 5
Segment 6
Segment 7
Segment 8
Segment 9
Segment 10
Segment 11
Segment 12
Density
Cosine Distance

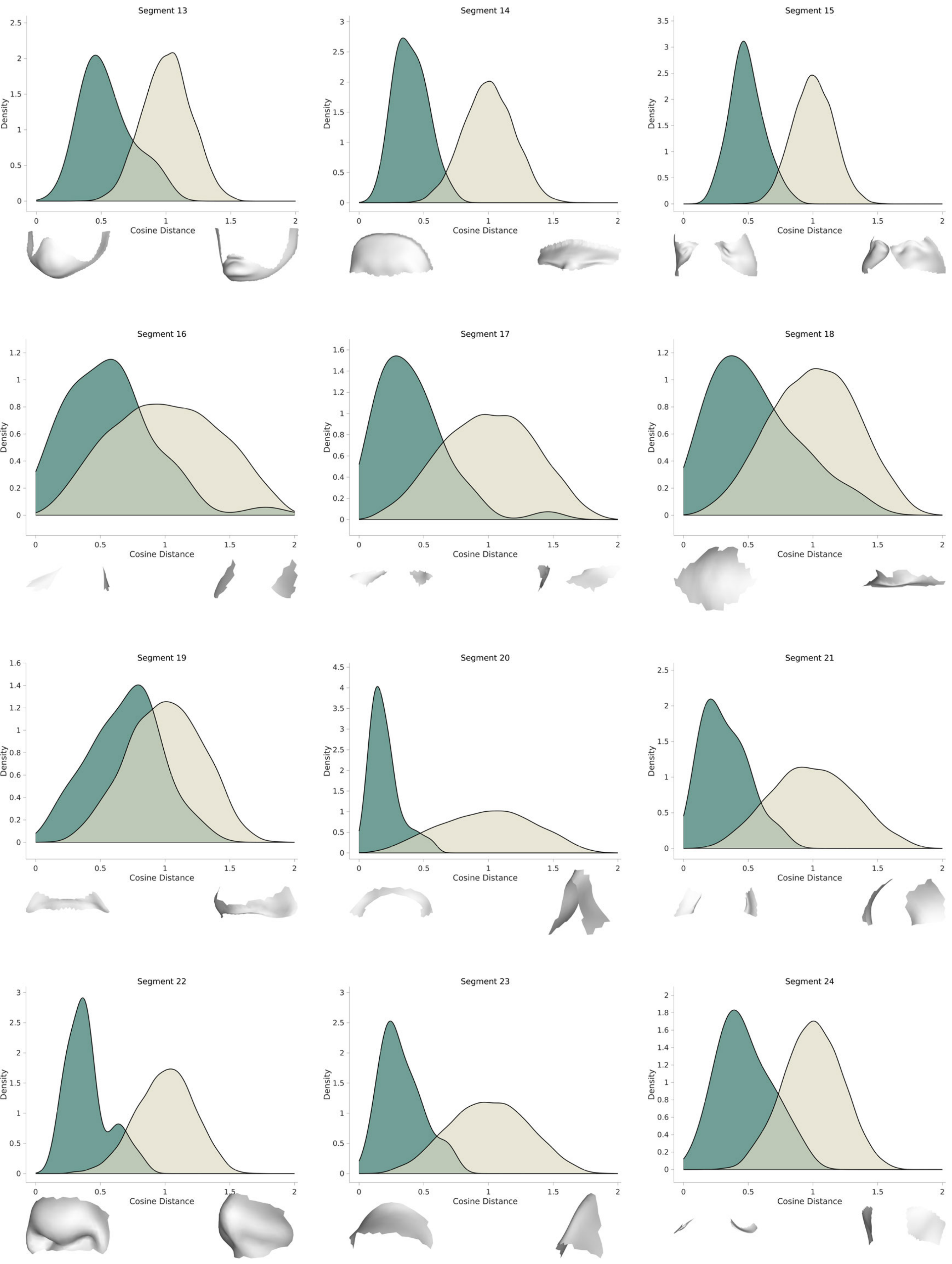
Segment 13
Segment 14
Segment 15
Segment 16
Segment 17
Segment 18
Segment 19
Segment 20
Segment 21
Segment 22
Segment 23
Segment 24
Density
Cosine Distance

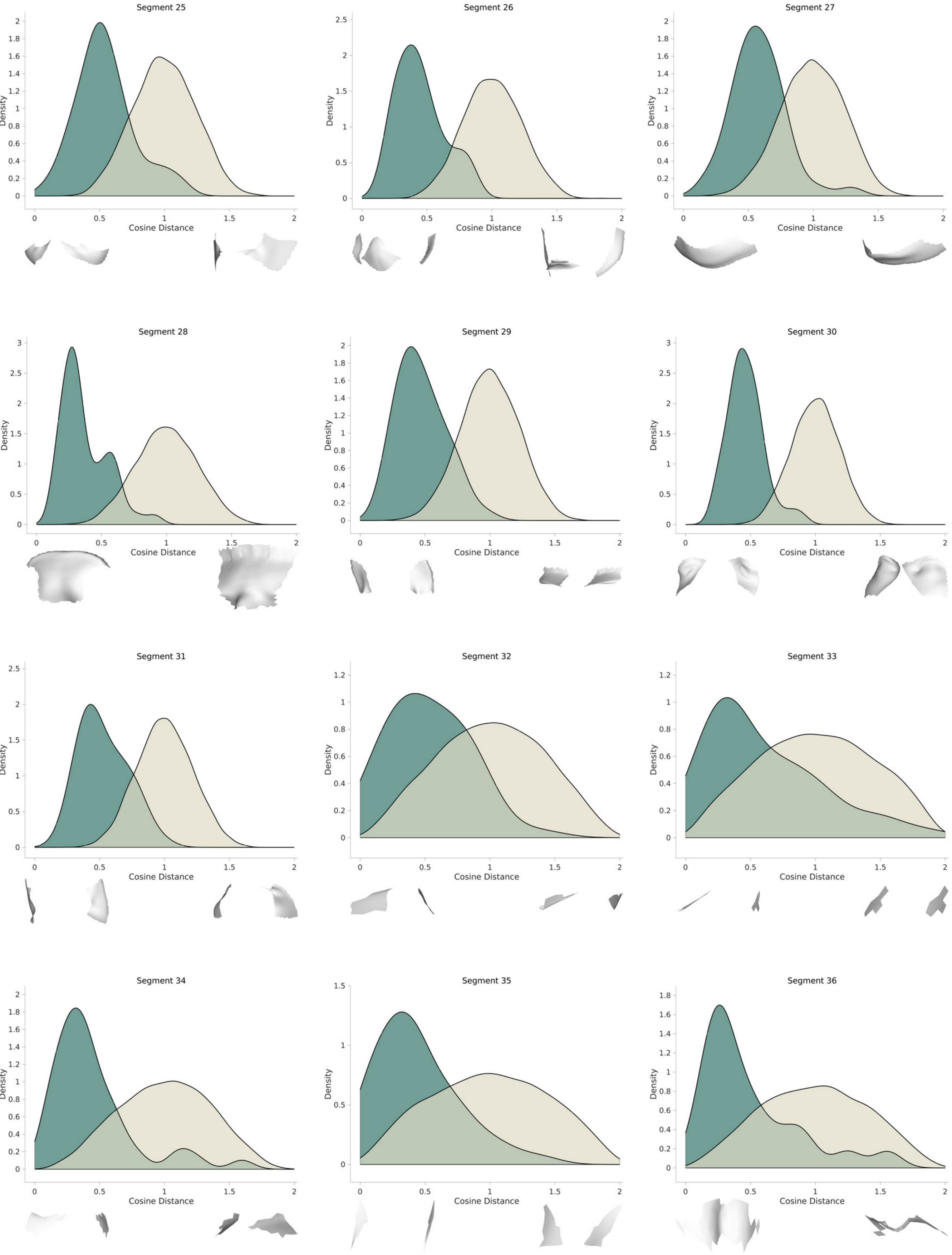
Segment 25
Segment 26
Segment 27
Segment 28
Segment 29
Segment 30
Segment 31
Segment 32
Segment 33
Segment 34
Segment 35
Segment 36
Density
Cosine Distance

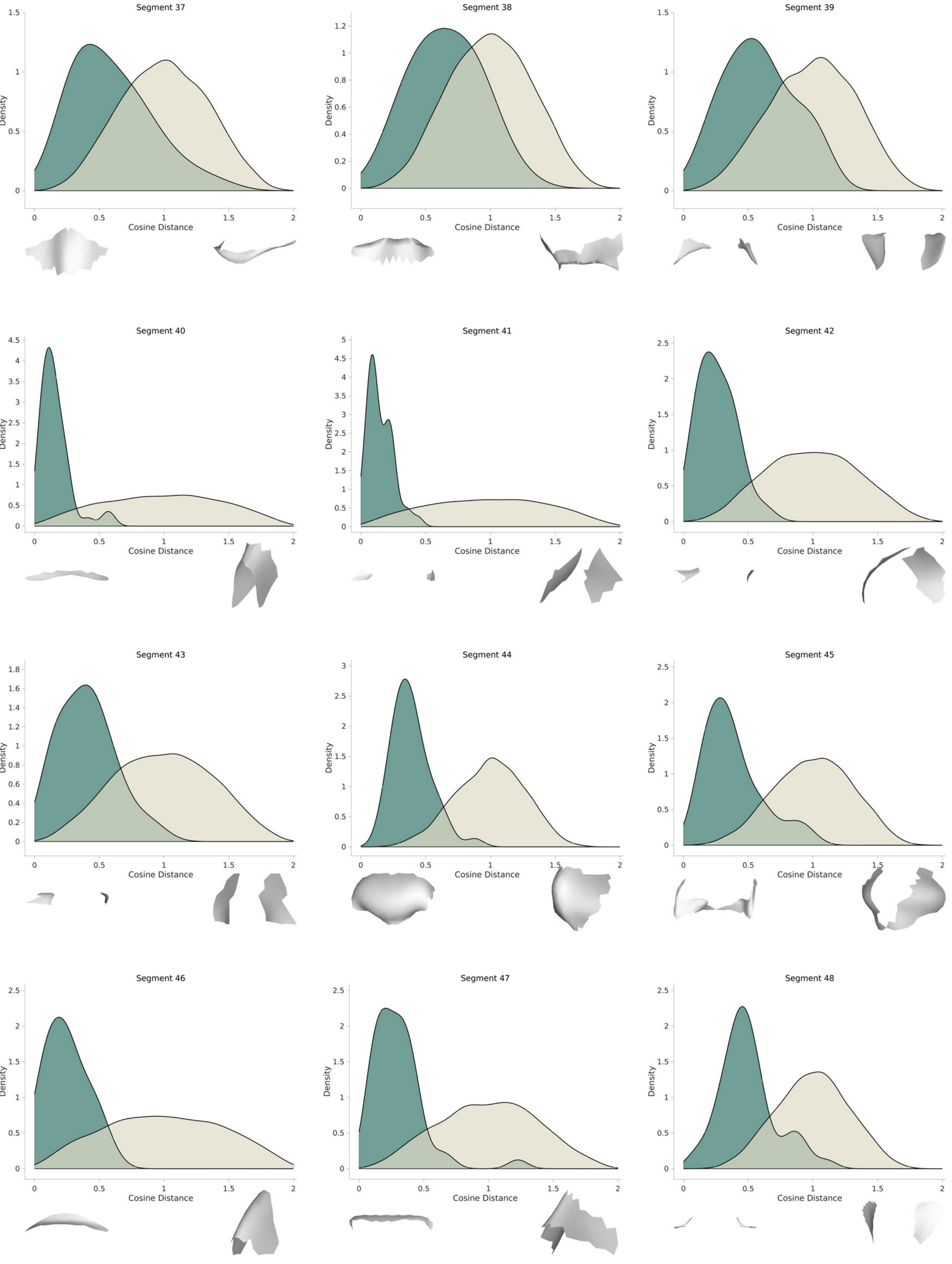

Segment 37
Segment 38
Segment 39
Segment 40
Segment 41
Segment 42
Segment 43
Segment 44
Segment 45
Segment 46
Segment 47
Segment 48
Density
Cosine Distance

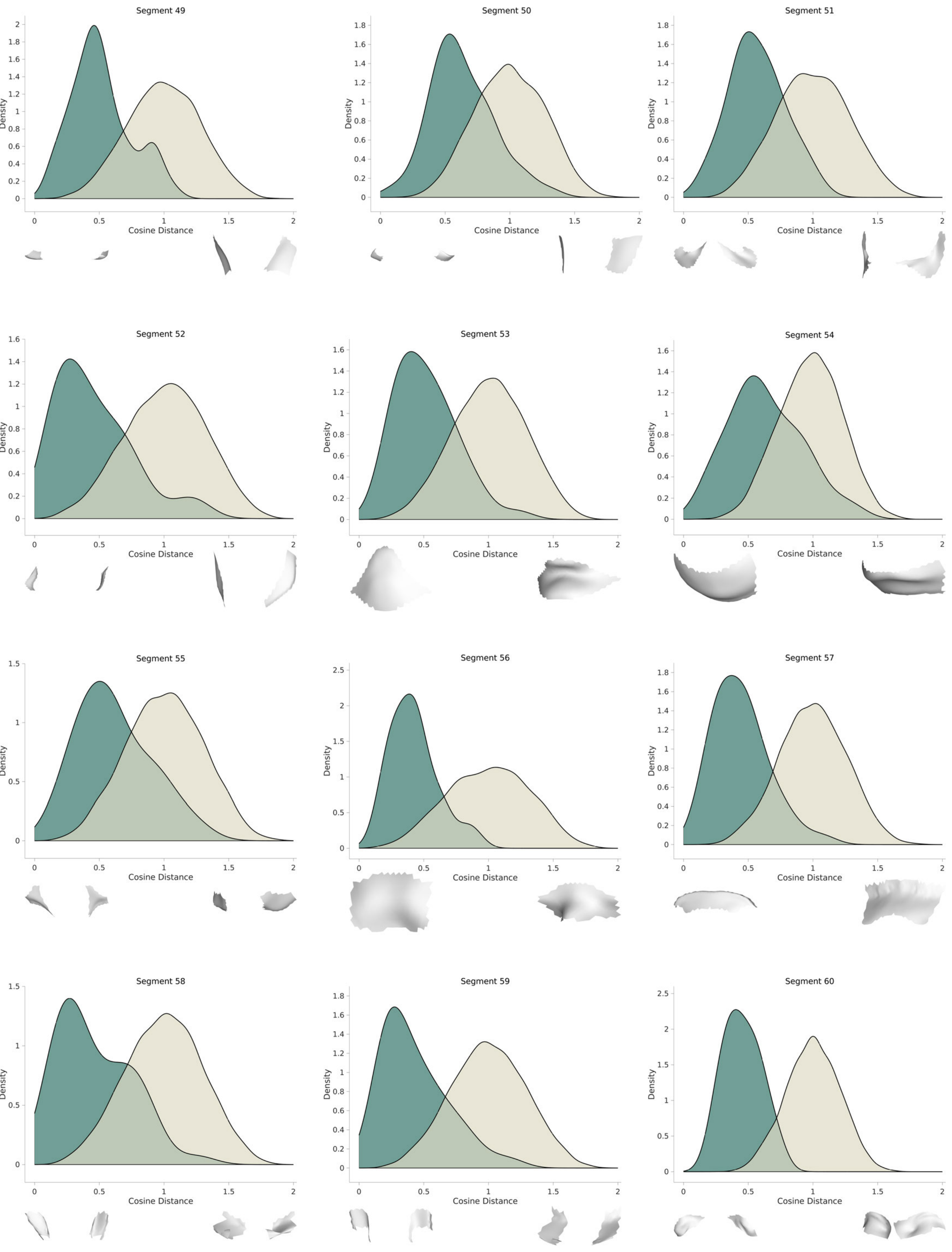

Segment 49
Segment 50
Segment 51
Segment 52
Segment 53
Segment 54
Segment 55
Segment 56
Segment 57
Segment 58
Segment 59
Segment 60
Density
Cosine Distance

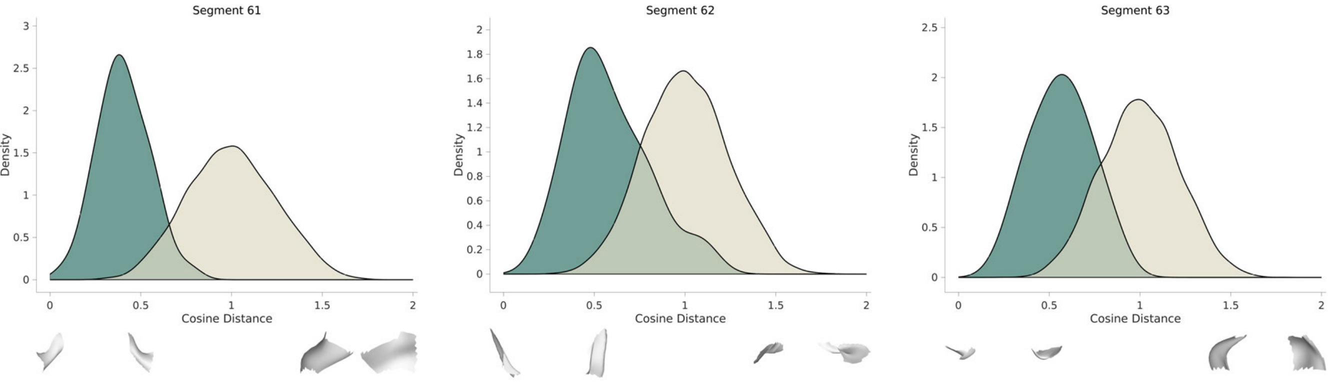
Segment 61
Segment 62
Segment 63
Density
Cosine Distance
0
0.5
1
1.5
2
2.5
3
0.2
0.4
0.6
0.8
1.2
1.4
1.6
1.8
