## Supplementary Data 3 for "Syndrome-informed phenotyping identifies a polygenic background for achondroplasia-like facial variation in the general population"

**Supplementary Data 3. Facial shape effects associated with the 35 lead SNPs.** The ACH trait scores obtained per facial segment were combined into a matrix, and a multivariate GWAS was performed using canonical correlation analysis (CCA). For each lead SNP, the dendrogram and facial heatmaps show the absolute values of the CCA loadings per segment. The position of the nodes corresponds to the facial segments depicted in Supplementary Figure 2a. Facial morphs are plotted separately for each hierarchical level.

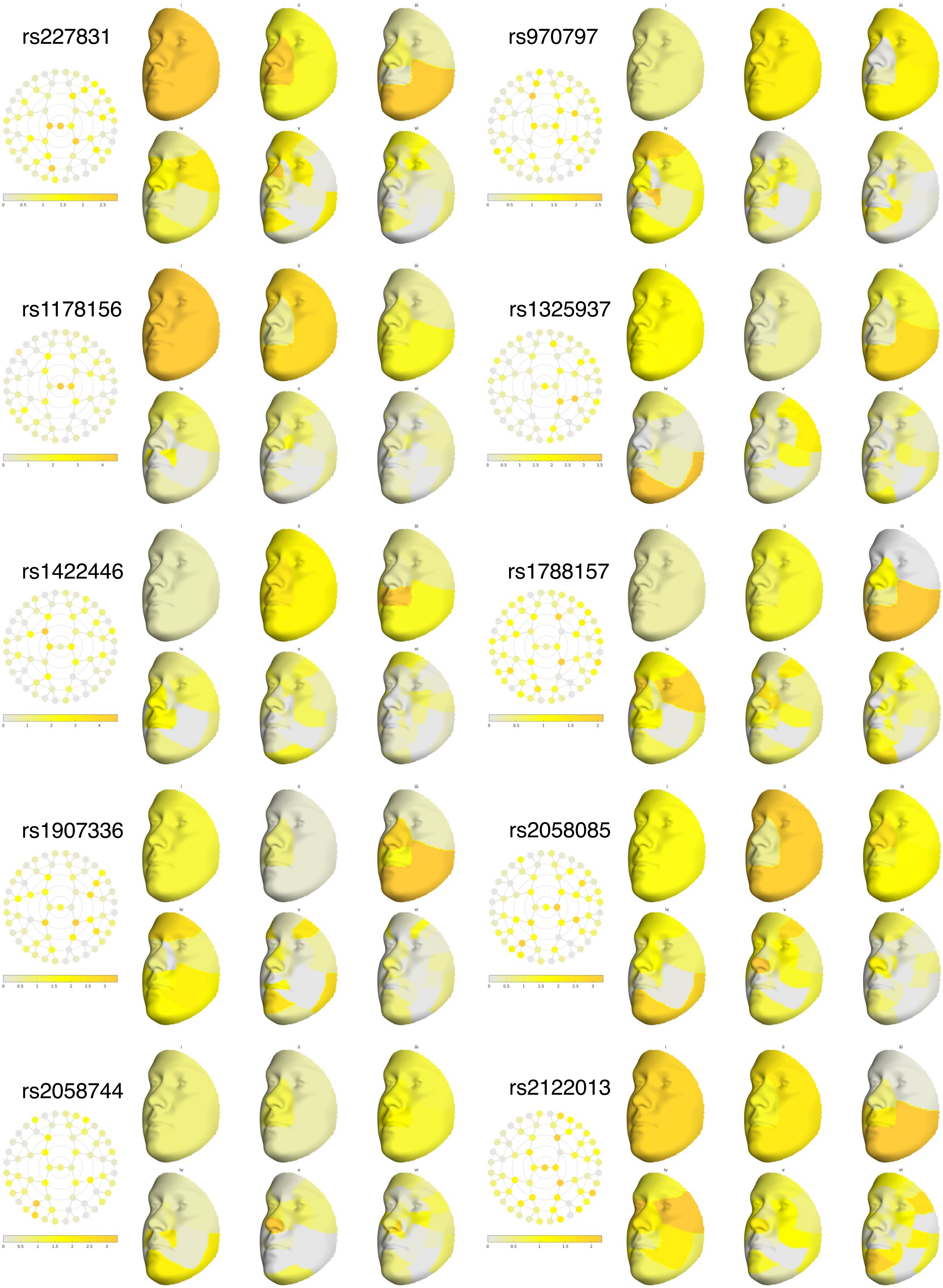
rs227831
rs970797
rs1178156
rs1325937
rs1422446
rs1788157
rs1907336
rs2058085
rs2058744
rs2122013

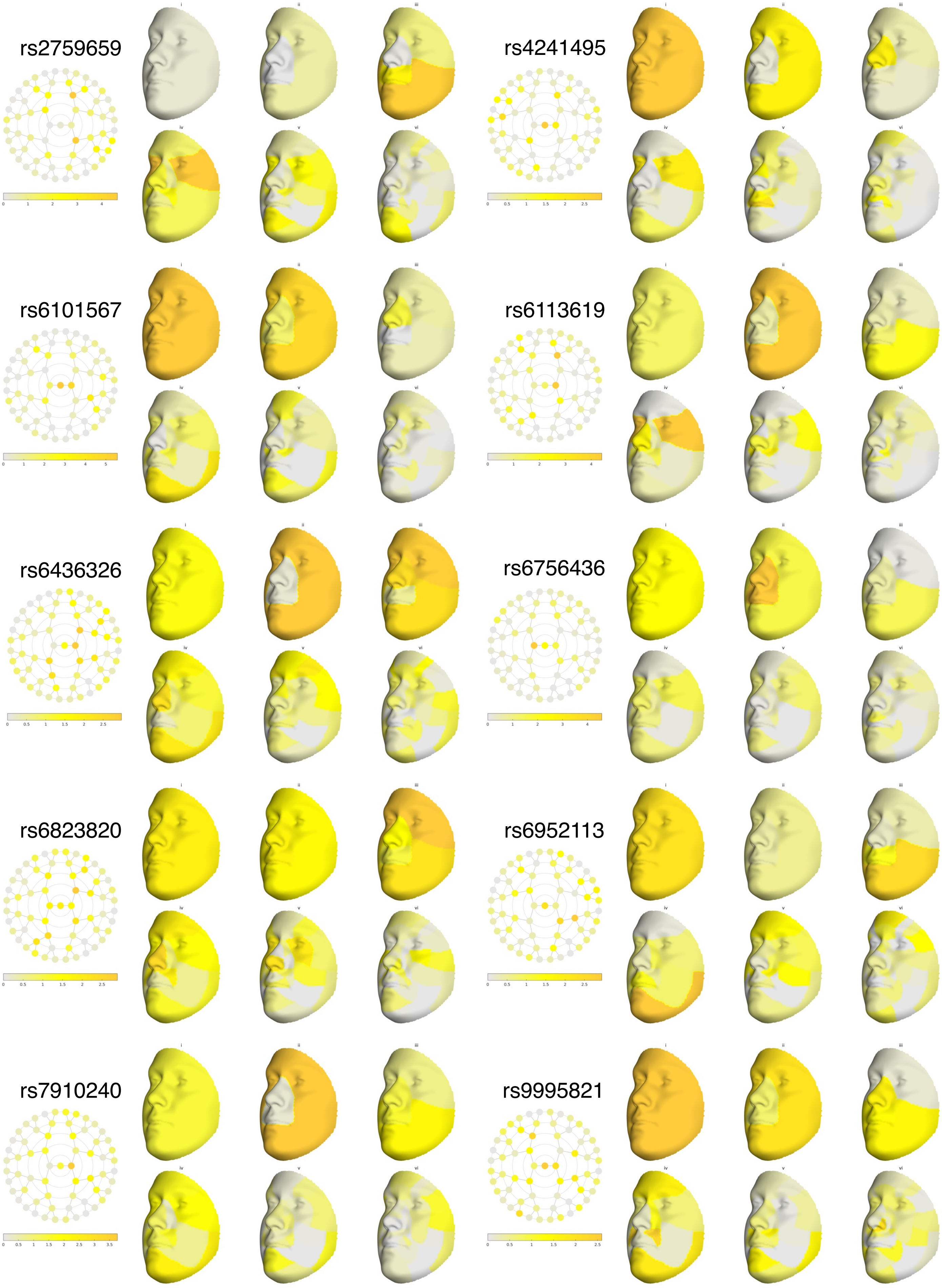
rs2759659
rs4241495
rs6101567
rs6113619
rs6436326
rs6756436
rs6823820
rs6952113
rs7910240
rs9995821

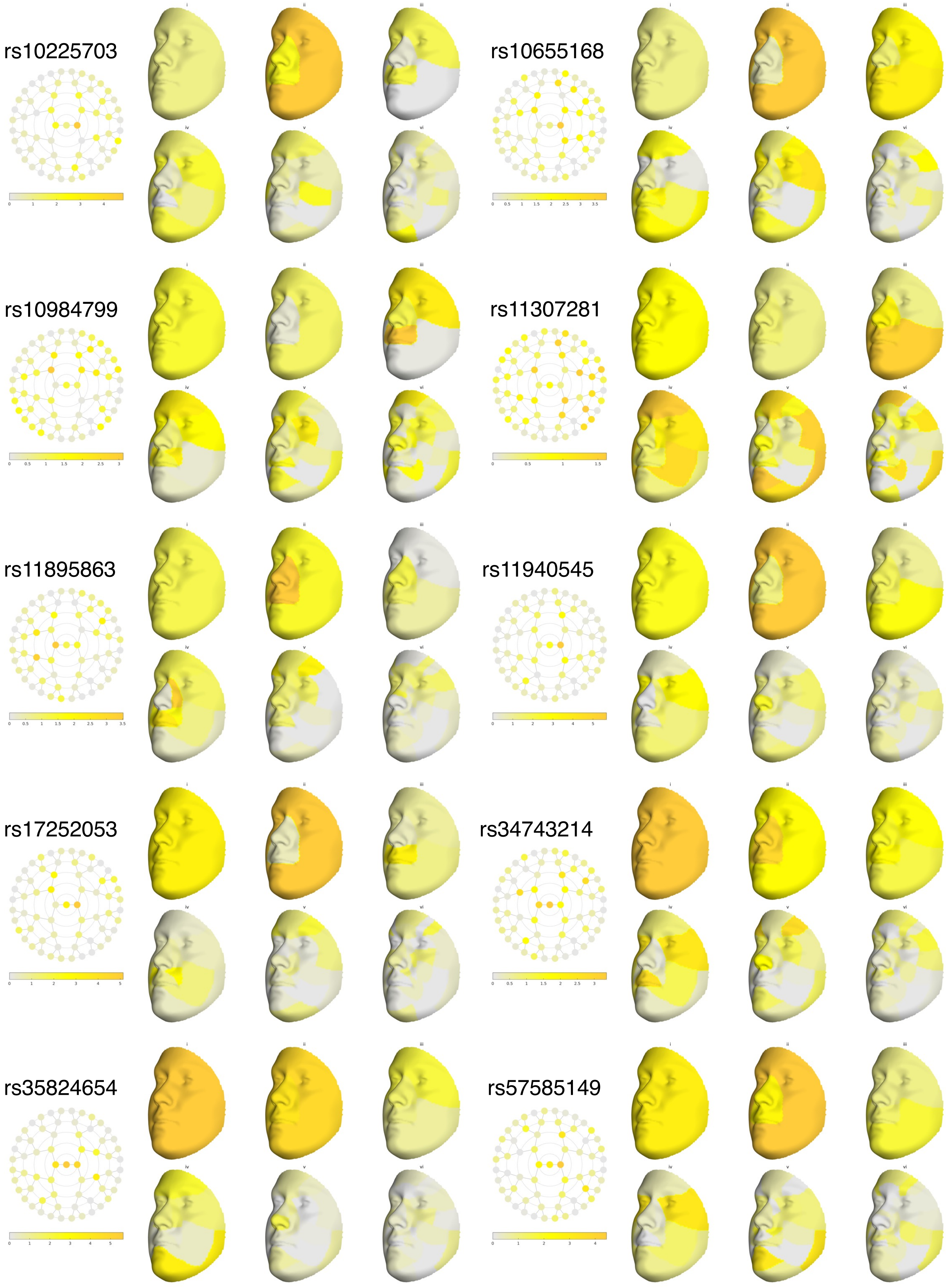
rs10225703
rs10655168
rs10984799
rs11307281
rs11895863
rs11940545
rs17252053
rs34743214
rs35824654
rs57585149

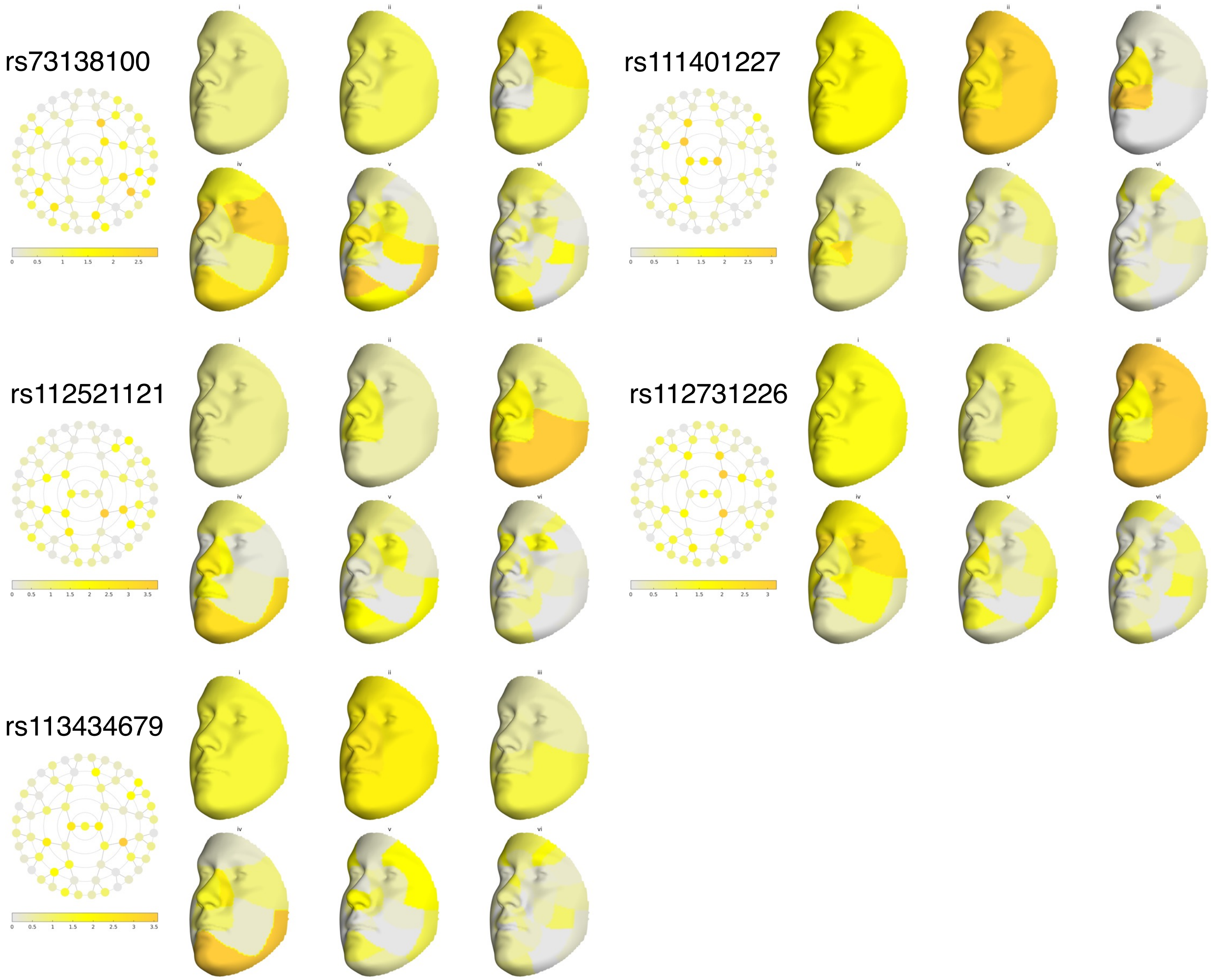
rs73138100
rs111401227
rs112521121
rs112731226
rs113434679
